# *in vivo* quantitative FRET small animal imaging: intensity versus lifetime-based FRET

**DOI:** 10.1101/2023.01.24.525411

**Authors:** Jason T. Smith, Nattawut Sinsuebphon, Alena Rudkouskaya, Xavier Michalet, Xavier Intes, Margarida Barroso

## Abstract

Förster Resonance Energy Transfer (FRET) microscopy is used in numerous biophysical and biomedical applications to monitor inter- and intramolecular interactions and conformational changes in the 2–10 nm range. FRET is currently being extended to *in vivo* optical imaging, its main application being in quantifying drug-target engagement or drug release in animal models of cancer using organic dye or nanoparticle-labeled probes. Herein, we compared FRET quantification using intensity-based FRET (sensitized emission FRET analysis with the 3-cube approach using an IVIS imager) and macroscopic fluorescence lifetime (MFLI) FRET using a custom system using a time-gated ICCD, for small animal optical *in vivo* imaging. The analytical expressions and experimental protocols required to quantify the product *f*_*D*_ *E* of the FRET efficiency *E* and the fraction of donor molecules involved in FRET, *f*_*D*_, are described in detail for both methodologies. Dynamic *in vivo* FRET quantification of transferrin receptor-transferrin binding was acquired in live intact nude mice upon intravenous injection of near infrared-labeled transferrin FRET pair and benchmarked against *in vitro* FRET using hybridized oligonucleotides. Even though both *in vivo* imaging techniques provided similar dynamic trends for receptor-ligand engagement, we demonstrate that MFLI FRET has significant advantages. Whereas the sensitized emission FRET approach using the IVIS imager required 9 measurements (6 of which are used for calibration) acquired from three mice, MFLI FRET needed only one measurement collected from a single mouse, although a control mouse might be needed in a more general situation. Based on our study, MFLI therefore represents the method of choice for longitudinal preclinical FRET studies such as that of targeted drug delivery in intact, live mice.

**WHY IT MATTERS:** FRET measurements in live animals open a unique window into drug-target interaction monitoring, by sensing the close proximity between a donor and acceptor-labeled molecular probes. To perform these measurements, a 3-cube fluorescent intensity measurement strategy can be adopted, as is common for *in vitro* FRET microscopy studies. However, it is challenging to translate this already cumbersome approach to *in vivo* small animal imaging. Here, we compare this standard approach, for which we provide a revised analytical framework, to a conceptually much simpler and more powerful one based on fluorescence lifetime measurements. Our results demonstrate that the technical challenge of *in vivo* fluorescence lifetime macroscopic imaging is well worth surmounting to obtain quantitative, whole-animal information regarding molecular drug-target engagement.

## 1 INTRODUCTION

Förster Resonance Energy Transfer (FRET) has been extensively used in fluorescence microscopy as a nanometer-range (2-10 nm) proximity assay (1, 2), addressing a distance range that even super-resolution microscopy cannot resolve (< 20-30 nm) in live cells. FRET provides information on the distance between donor (D) labeled and acceptor (A) labeled proteins for each specific donor-acceptor fluorophore pair, independently of the resolution provided by the fluorescence imaging methodology used to acquire FRET measurements (3–6). Thus, FRET can be performed at both visible as well as near-infrared (NIR) wavelengths and can be measured by a wide variety of fluorescence-based imaging methodologies, beyond traditional microscopy approaches (7–9). These characteristics make FRET broadly applicable and one of the most extensively used imaging approaches in living cells as well as in model organisms, including bacteria, yeast, C. *elegans*, drosophila and mice (6).

There are several different types of FRET assays in fluorescence biological imaging. *Intra*-molecular FRET is used mostly to detect transient and dynamic signaling events using genetically encoded FRET-based biosensors in living cells (10–14). These biosensor constructs provide a constant 1:1 donor:acceptor stoichiometry in each pixel, allowing for the implementation of ratiometric intensity-based imaging for a fast and qualitative FRET analysis. However, ratiometric FRET is very sensitive to the signal-to-noise ratio (SNR), has limited dynamic range and requires a significant number of image processing steps, such as background subtraction, shade/flatfield correction, image alignment and photobleaching correction as discussed in the literature(15–25). These problems are further compounded for visible range fluorophores in tissues, where measurements are affected by autofluorescence and the wavelength and tissue-dependent attenuation of light propagation in heterogeneous tissues (26). *Inter*-molecular FRET has been established to monitor protein-protein interactions in live cells using separate donor- and acceptor-labeled proteins (27, 28). However, in inter-molecular FRET the relative abundance of donor and acceptor fluorophores is not always controllable and can change over time, limiting the information that can be extracted from ratiometric measurements to the *apparent* or *average* energy transfer efficiency ⟨*E*⟩, which depends on the D to A distance (FRET efficiency *E* for D-A pairs) but also on the fraction *f*_*D*_ of donor molecules that are involved in energy transfer.

Fluorescence lifetime microscopy (FLIM) is regarded as the most robust means to collect FRET data since it is largely not influenced by probe concentration, signal intensity, or spectral bleed-through contamination (6). FLIM quantifies FRET occurrence by measuring the reduction of the fluorescence lifetime of the donor when in close proximity to the acceptor. Since the acceptor effectively behaves as a quencher of the donor’s fluorescence, this quenching process is accompanied by a reduction in the quantum yield and lifetime of the donor (7). FLIM can measure FRET in each biological sample via the collection of the donor emission channel only. However, FLIM requires complex instrumentation and fairly advanced analysis involving either the phasor approach or model-based fitting (29), which makes it less accessible or straightforward compared to intensity-based FRET, which can be implemented with standard fluorescence microscopes and involves simple algebraic data processing (30–32). On the other hand, FLIM-FRET is not devoid of potential traps, as a fluorophore’s lifetime can be sensitive to many other environmental perturbations (33), and should therefore be carried with appropriate control experiments.

Extending FRET assays to *in vivo* non-invasive *macroscopy* is one of the last frontiers of FRET imaging. Recently, *in vivo* FRET imaging approaches have been implemented to measure nanoparticle drug delivery and release, drug-target engagement, and dynamic probe uptake or biosensor-based signaling in various pre-clinical animal models (8, 9, 34–39). A major issue preventing full application of FRET into small animal imaging is the need to red-shift FRET into the NIR range to reduce absorption and minimize autofluorescence, as well as to increase depth of penetration in thick tissues (40). Development of NIR-labeled donor and acceptor pairs has permitted the implementation of non-invasive longitudinal FRET as well as the multiplexing of FRET pairs with metabolic imaging application in intact living mice (41–44), although that comes with additional challenges, such as the shorter fluorescence lifetime and lower quantum yield of NIR emitting dyes.

Here, we address this challenge by comparing intensity- and lifetime-based NIR intermolecular FRET imaging assays designed to monitor receptor-ligand interactions in live intact mice (41, 44–47). In the context of ligand-receptor systems, FRET between donor-labeled and acceptor-labeled ligands occurs upon their binding to membrane-bound dimerized receptors. Using intensity- and lifetime-based FRET microscopy, we have demonstrated that protein ligands (*e. g*. transferrin: Tf), do bind extracellular domains of membrane-bound receptors (*e. g*. transferrin receptor: TfR) (48, 49). Moreover, *in vivo* MFLI FRET measurements have been successfully validated via *ex vivo* histochemistry, establishing that *in vivo* FRET signal directly reports on receptor-ligand engagement in intact live animals (43, 44, 47, 50, 51).

In the present study, we revisited the standard 3-cube equations for intensity-based FRET in the NIR range and systematically compared its results to lifetime-based FRET measurements analysis for macroscopic imaging. The comparison was first done *in vitro* with NIR-labeled double-stranded DNA FRET standard samples. We then extended our comparison to *in vivo* pharmaco-kinetics of NIR-labeled ligand-receptor engagement monitored over more than one hour and a half. Altogether, we show that while intensity-based NIR FRET analysis *in vivo* can be performed, lifetime-based *in vivo* NIR FRET analysis is a much more robust and reliable approach for whole-animal quantitative FRET imaging.

## 2 MATERIAL AND METHODS

### 2.1 Macroscopic fluorescence lifetime-FRET (MFLI-FRET) with gated-ICCD

MFLI was performed using a time-resolved wide-field illumination and a a time-gated intensified charge-coupled device (ICCD) camera (42). The system’s excitation source was a tunable fs pulsed Ti-Sapphire laser (Mai Tai HP, Spectra-Physics, CA) set to 695 nm. Power at the imaging plane was approximately 2 and 3 mW/cm^2^ for *in vitro* and *in vivo* MFLI, respectively. A digital micro-mirror device (DLi 4110, Texas Instruments, TX) was used for wide-field illumination over the sample plane. During animal imaging, active illumination was applied to ensure that the signal in the different regions of interest did not saturate the camera (52) (Supplemental Table S1). The time-gated ICCD camera (Picostar HR, LaVision, Germany) was set to acquire gate images with a gate width of *W*_*ICCD*_ = 300 ps, separated by gate steps *δt* = 40 ps (details provided in ref. (42)). These width and resolution have been shown to be sufficient for efficient recovery of the short lifetimes involved in this study (and in fact wider and sparser IRF measurements have been successfully used as well (51)), provided a sufficient SNR is achieved. Gates covered over a temporal window of duration shorter than the full 12.5 ns laser period (*G* = 150 to 176 total gate images per acquisition, *i.e. D* = 6 to 7 ns), sufficient to acquire the full fluorescence decay. During fluorescence imaging, a bandpass filter 720 ± 6.5 nm (FF 720/13, Semrock, NY) and a longpass filter 715 nm (FF 715/LP25, Semrock, NY) were applied to selectively collect donor fluorescence signal and reject laser scatter and acceptor fluorescence. The ICCD’s microchannel plate (MCP) voltage and the gate integration time were further optimized in each case to avoid detector saturation (Supplemental Table S1). Instrument response functions (IRFs) were acquired with equivalent illumination conditions to those used for fluorescence imaging, except for the emission filters, which were removed.

### 2.2 Intensity-based FRET imaging using IVIS Imager

All samples, including donor-only (DO), acceptor-only (AO) and double-labeled (DA) samples, were imaged simultaneously in the same field-of-view of an IVIS Lumina XRMS Series III imaging system (Perkin Elmer, MA), including heated stage (37 °C) and isoflurane anesthesia connections for small animal imaging. Excitation wavelengths were set to 660 ± 10 nm for the donor and 740 ± 10 nm for the acceptor fluorophores. The emission filters were set to 713 ± 20 nm for the donor and to 793 ± 20 nm for the acceptor fluorophores. The intensity used in IVIS was constant throughout all imaging experiments. Three spectral channels were acquired for intensity FRET imaging: 1) donor channel (donor excitation and donor emission), 2) acceptor channel (acceptor excitation and acceptor emission) and 3) FRET channel (donor excitation and acceptor emission), with adjusted exposure time for each channel (Supplemental Table S2). The image size was 256×256 pixels after 4×4 binning of the camera full-frame image. In the case of *in vivo* imaging, one set of images (donor, acceptor and FRET channels) was acquired before any fluorophore injection and used as background and subtracted from the subsequent series.

### 2.3 Intensity-based FRET Data Analysis

Intensity-based FRET efficiency measurement relies on quantifying the amount of FRET-induced acceptor fluorescence (also called sensitized emission) in a sample, relative to that measured in the donor emission channel.

In an ideal situation where each donor fluorophore is located at a fixed distance from an acceptor fluorophore, (*i. e*. samples in which 100% of the donor molecules undergo FRET, denoted DA to emphasize that each donor forms a pair with an acceptor), a simple ratiometric approach using only signals obtained upon excitation with a donor-specific wavelength can be used to obtain the so-called proximity ratio (PR), or uncorrected ratiometric FRET efficiency, given by Eq. 1 (53):

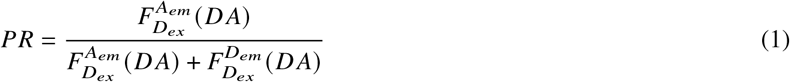

where 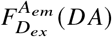 and 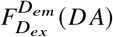 are background-corrected acceptor and donor intensities of the sample undergoing FRET measured upon donor excitation, respectively (see Table 1 for notations). Measuring PR is a semi-quantitative approach for approximately quantifying FRET efficiency in a FRET sample where donor and acceptor are for instance conjugated to the same molecule, but it leaves aside contributions such as donor emission crosstalk (donor signal detected in the acceptor emission channel) and acceptor cross-excitation (direct excitation of the acceptor with donor excitation wavelengths) among other effects. Indeed, generally, the total fluorescence collected in each emission channel is a contribution of acceptor emission from FRET, donor emission leakage, and acceptor emission from direct excitation.

**Table 1:**
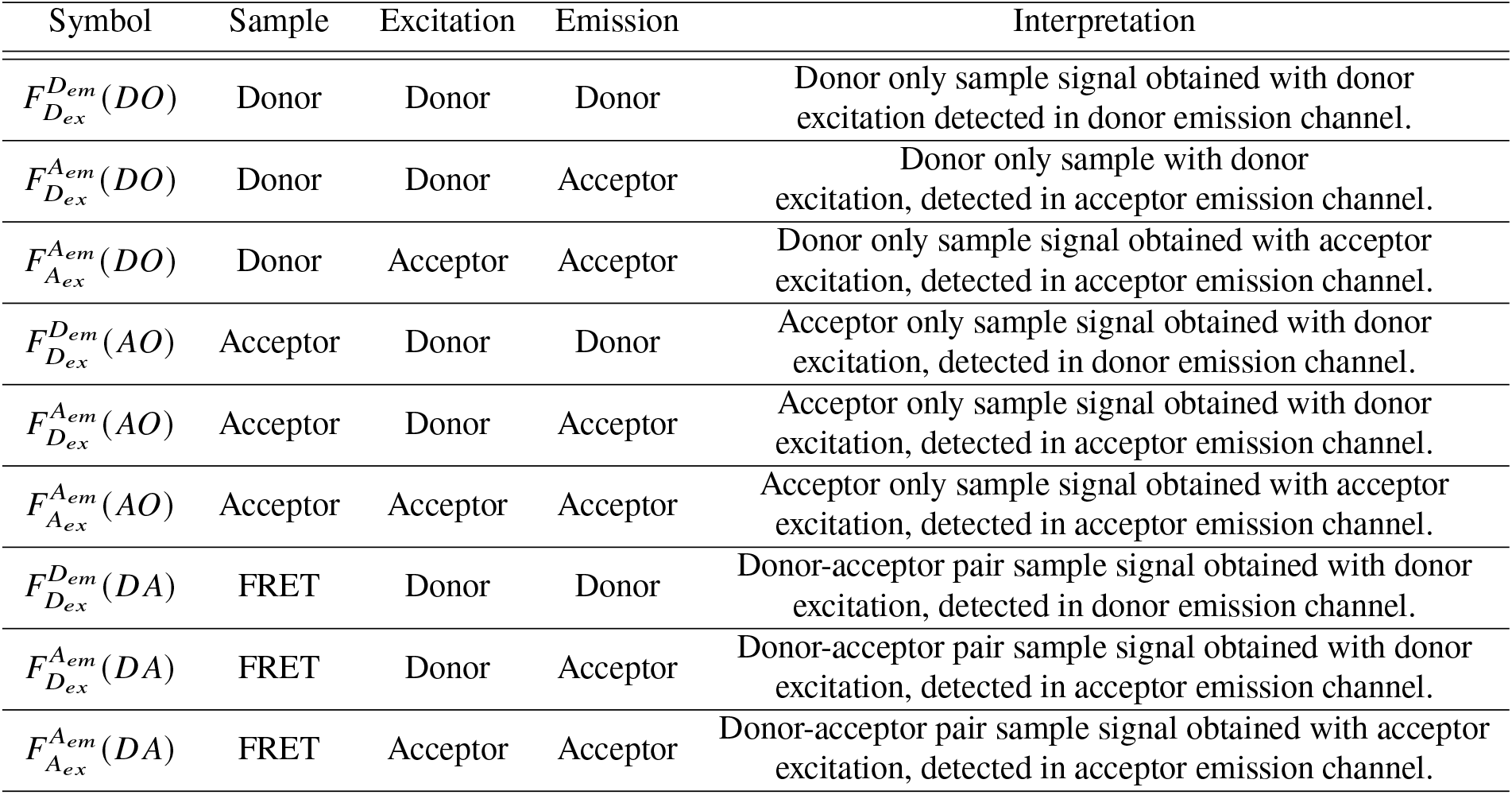
Notations used in the text to refer to various background-corrected sample signals and their description. The notation 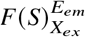, used in ref. (53), represents the signal from species “*S*” (e.g. donor *D*) excited by excitation channel *X* (lower index notation, e.g. *D*_*ex*_ for donor excitation laser) and detected in emission channel *E* (e.g. *D*_*em*_ for donor emission channel.

| Symbol | Sample | Excitation | Emission | Interpretation |
| --- | --- | --- | --- | --- |
| $F_{D_{ex}}^{D_{em}}(DO)$ | Donor | Donor | Donor | Donor only sample signal obtained with donor excitation detected in donor emission channel. |
| $F_{D_{ex}}^{A_{em}}(DO)$ | Donor | Donor | Acceptor | Donor only sample with donor excitation, detected in acceptor emission channel. |
| $F_{A_{ex}}^{A_{em}}(DO)$ | Donor | Acceptor | Acceptor | Donor only sample signal obtained with acceptor excitation, detected in acceptor emission channel. |
| $F_{D_{ex}}^{D_{em}}(AO)$ | Acceptor | Donor | Donor | Acceptor only sample signal obtained with donor excitation, detected in donor emission channel. |
| $F_{D_{ex}}^{A_{em}}(AO)$ | Acceptor | Donor | Acceptor | Acceptor only sample signal obtained with donor excitation, detected in acceptor emission channel. |
| $F_{A_{ex}}^{A_{em}}(AO)$ | Acceptor | Acceptor | Acceptor | Acceptor only sample signal obtained with acceptor excitation, detected in acceptor emission channel. |
| $F_{D_{ex}}^{D_{em}}(DA)$ | FRET | Donor | Donor | Donor-acceptor pair sample signal obtained with donor excitation, detected in donor emission channel. |
| $F_{D_{ex}}^{A_{em}}(DA)$ | FRET | Donor | Acceptor | Donor-acceptor pair sample signal obtained with donor excitation, detected in acceptor emission channel. |
| $F_{A_{ex}}^{A_{em}}(DA)$ | FRET | Acceptor | Acceptor | Donor-acceptor pair sample signal obtained with acceptor excitation, detected in acceptor emission channel. |

Sensitized emission FRET (SE-FRET) approaches have been designed to correct for these additional effects and require data acquired with separate excitation and emission combinations (the so-called 3-cube approach) (16, 22, 53–57).

A first-order correction consists in subtracting the direct acceptor excitation and the leakage of the donor emission from the measured acceptor signal to obtain a better estimate of the FRET-induced acceptor emission (*i.e*., the relevant FRET emission signal), using Eq. 2:

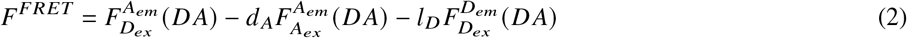

where *d* _*A*_ is the direct acceptor excitation correction factor, and *l*_*D*_ is the donor leakage correction factor.

The first correction factor *d* _*A*_ is measured using an acceptor-only (AO) sample excited separately at two excitation wavelengths (donor and acceptor) and detected in the acceptor emission channel. Correction factor *d* _*A*_ is calculated using Eq. A.13 in the Appendix (17, 24, 53, 57).

The second correction factor *l*_*D*_ is measured using a donor-only (DO) sample excited with a donor excitation wavelength (donor excitation channel) and detected in both emission channels (donor and acceptor). Correction factor *l*_*D*_ is calculated using Eq. A.12 in the Appendix (17, 24, 53, 57).

The FRET efficiency *E* can then be computed as (24, 53):

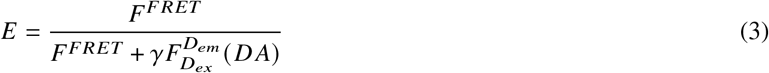

where *γ* is the detection-correction factor defined as 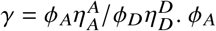. *ϕ*_*A*_ and *ϕ*_*D*_ are the acceptor and donor quantum yields, respectively and 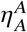(resp.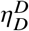) is the acceptor (resp. donor) detection efficiency in the acceptor (resp. donor) channel.

While Eqs. 2 and 3 are adequate in many situations, certain experimental situations result in further signal contamination, when for instance the donor fluorophore can be excited at the acceptor excitation wavelength, or when the acceptor fluorophore can be detected in the donor emission channel. The first effect contributes some unwanted signal to a quantity used to correct the sensitized emission of the acceptor in Eq. 2, while the second requires further correction of the donor channel signal. In those cases, some donor signal needs to be subtracted from 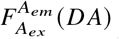 and some acceptor signal from 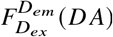.

These corrections involve two additional correction factors (*d*_*D*_ and *l*_*A*_), as discussed in the Appendix (54). To retrieve *d*_*D*_, two measurements of a donor-only sample are needed: i) excitation at the donor wavelength and recording in the acceptor emission channel and ii) excitation at the acceptor wavelength and recording in the acceptor emission channel. Correction factor *d*_*D*_ is calculated using Eq. A.12 in the Appendix (54).

The last correction factor *l*_*A*_ requires two measurements of an acceptor-only sample: i) excitation at the donor wavelength and recording in the donor emission channel and ii) excitation at the donor wavelength and recording in the acceptor emission channel. Correction factor *l* _*A*_ is calculated using Eq. A.13 in the Appendix (54).

It should be noted that correction factors *d* _*A*_, *l* _*A*_, *d*_*D*_ and *l*_*D*_ are specific to fluorophores as well as imaging systems, and ideally require constant excitation intensities throughout the series of measurements. At the very least, one must take into account differences in excitation intensity (and integration time) if those need to be adjusted for experimental reasons. Consequently, these correction factors need to be estimated every time the experimental conditions are modified (excitation intensities, integration times, filters, fluorophores or molecular environments).

After all four correction terms have been retrieved, the sensitized emission FRET signal, *F*^*FRET*^, can be calculated according to (see Appendix Eq. A.21 for details):

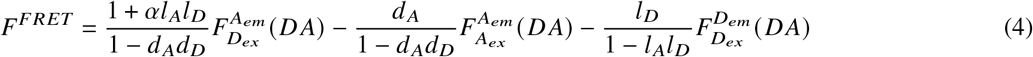

where parameter *α* is defined by Eq. A.19 in the Appendix. If *l* _*A*_ = *d*_*D*_ = 0, one recovers Eq. 2.

Similarly, the FRET efficiency can be obtained by a modified version of Eq. 3 (see Appendix Eq. A.22):

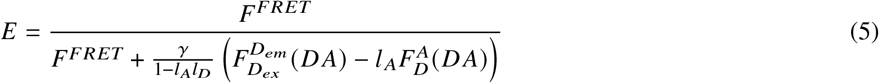

an expanded version of which can be found in the Appendix (Eq. A.18). If *l* _*A*_ = *d*_*D*_ = 0, one obviously recovers Eq. 3.

The above equations only apply to a pure FRET sample (DA), as mentioned at the beginning of the discussion and indicated by the notations. In general, a real sample will contain a mixture *M* of species: donor-only (DO), acceptor-only (AO) and FRET (DA), whose respective fractions are fully specified by the total number *N*_*D*_ of donor molecules and total number *N*_*A*_ of acceptor molecules, and the fraction *f*_*D*_ of donor molecules and fraction *f*_*A*_ of acceptor molecules involved in FRET interaction (with *f*_*D*_ *N*_*D*_ = *f*_*A*_*N*_*A*_). This mixture of species will be characterized by 3 different types of signals 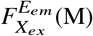, each verifying:

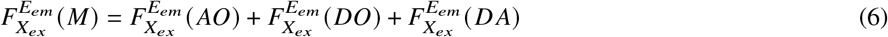

As derived in the Appendix, the product *f*_*D*_ *E* of the fraction of donor *f*_*D*_ involved in FRET and the FRET efficiency *E* of the FRET sample can then be expressed in terms of the 3 measured quantities 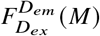, 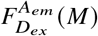 and 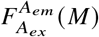 and the coefficients defined by Eqs. A.12-A.13,A.15 & A.19 as (Appendix Eq. A.27):

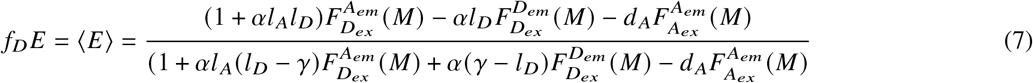

which turns out to be the same formula as obtained for a pure DA sample (Eq. 5) with the replacement of *E* by *f*_*D*_ *E*.

In the more general case where a number *n* of distinct FRET configurations {*D A*_*i*_}_*i*=1…*n*_ of the donor and acceptor molecules can be observed, with FRET efficiencies {*E*_*i*_} and fractions {*f*_*i*_}, the same formula applies, with the difference that the term *f*_*D*_ *E* is replaced by the sum 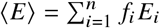 as shown in the Appendix (Eq. A.32).

Note in particular that, in the case of a mixture of D, A and a single DA species, it is not possible to disentangle *E* from *f*_*D*_ without further information on the sample. Fortunately, the quantity ⟨*E*⟩ can also be estimated using lifetime measurements as discussed in the next section, allowing a direct comparison of both methods.

### 2.4 Lifetime-based FRET Data Analysis

In the ideal case, quantification of FRET using fluorescence lifetime imaging (FLI) only requires measuring the fluorescence lifetime of the donor undergoing FRET and that of the isolated donor (no FRET condition). The result of FRET is a reduction (quenching) of the donor fluorescence lifetime.

There are two conventional methods to obtain lifetime-based FRET quantification: 1) multi-exponential fitting and 2) phasor analysis (58, 59). We have demonstrated the equivalence of the two methods for *in vitro* and *in vivo* MFLI-FRET analysis in recent publications (46, 51), and will therefore not discuss the latter method any further. In the simplest FRET-FLI analysis case, two donor lifetimes contribute to the observed decay: τ_*DA*_ is the lifetime of the donor undergoing FRET and τ_*DO*_ is the lifetime of the donor not undergoing FRET. The resulting decay can therefore be modeled using a bi-exponential function (Eq. 8):

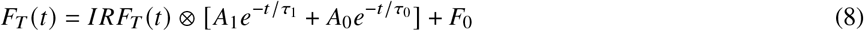

where *F*_*T*_ (*t*) is the *T*-periodic fluorescence intensity as function of time *t* after laser excitation (sometimes referred to as a temporal point spread function or TPSF). *IRF*_*T*_ (*t*) corresponds to the *T*-periodic instrument response function, which is convolved with the fluorescence decay (symbol ⊗, interpreted as a cyclic-convolution over a single period *T*) (60). *A*_1_ and *A*_0_ correspond to the amplitudes of the quenched and unquenched donor contributions, while τ_*DA*_ = τ_1_ and τ_*DO*_ = τ_0_ are the quenched and unquenched lifetimes, respectively.

The relative amplitudes of each component are related to the fraction of the donor in each species (donor-only and FRET pair) by (46):

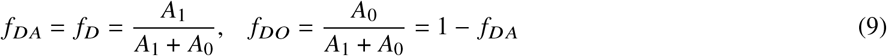

The unquenched donor lifetime (τ_*DO*_ = τ_0_) can be obtained experimentally (for instance as the longest lifetime component in a 2-exponential fit, or by a separate measurement of a donor-only sample acquired in identical conditions as the FRET sample) or from the literature. The *amplitude-weighted* average lifetime of the sample is calculated using Eq. 10, which is sometimes used as a “proxy” to quantify the fraction of FRET-undergoing species at a given location.

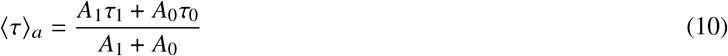

The donor-only lifetime (τ_*DO*_ = τ_0_) and the FRET sample lifetime (τ_*DA*_ = τ_1_) are related to the FRET efficiency (E) by:

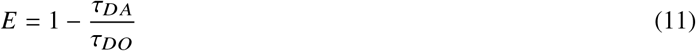

Combining Eqs. 9-11, we get the following expression for the product *f*_*D*_ *E*:

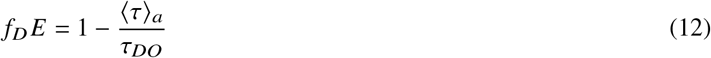

Because the FRET efficiency of the DO species is equal to zero (*E*_*DO*_ = 0), Eq. 12 can be rewritten as:

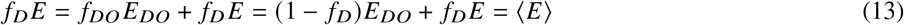

which states that the quantity *f*_*D*_*E* computed with Eq. 12 is the average FRET efficiency of the sample.

In the more general case where a number *n* of distinct FRET configurations *D A*_*i*_, *i* = 1…*n* of the donor and acceptor molecules can be observed, the lifetime τ_*DA*_ is replaced by *n* lifetimes {τ_*i*_}_*i*=1…*n*_, and the fraction *f*_*DA*_ by *n* fractions { *f*_*i*_}_*i*=1…*n*_ such that:

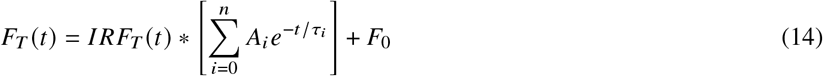

and:

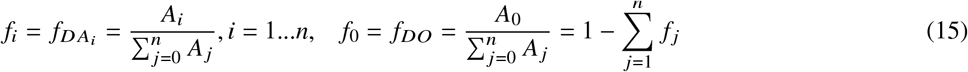

The *amplitude-weighted* average lifetime ⟨τ⟩_*a*_:

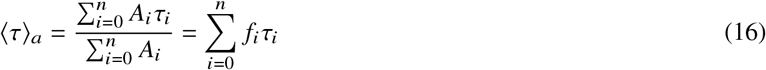

is therefore related to the mean FRET efficiency ⟨*E*⟩:

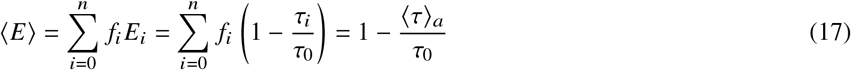

as before (τ_0_ = τ_*DO*_).

It is noteworthy that this expression has a simple interpretation in terms of the *average* FRET efficiency of the FRET-undergoing pairs. Indeed, defining the *amplitude-averaged* DA lifetime by:

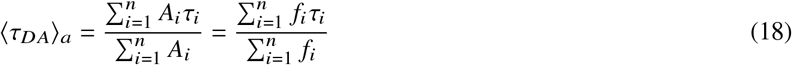

and the average FRET efficiency of the FRET-undergoing pairs as:

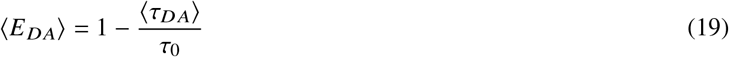

it follows that:

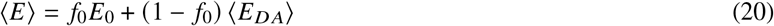

where *E*_0_ = *E*_*DO*_ = 0 is the donor-only FRET efficiency, showing that the occurrence of many different FRET configurations can be treated formally as equivalent to a situation with a single “average” configuration characterized by a FRET efficiency given by Eq. 19 (and a fraction given by 1 − *f*_0_). In practice, it is unlikely that a multi-exponential fit can be performed reliably enough to estimate the different (*A*_*i*_, τ_*i*_)’s needed to compute ⟨τ_*DA*_⟩_*a*_.

Eqs. 7 and 12 (or Eqs. A.32 and 17) provide a way to directly compare intensity-based and lifetime-based measurements of the same sample.

Importantly, acquisition of donor fluorescence lifetime data for MFLI-FRET quantification does not require acceptor fluorescence information (either acceptor emission recording or acceptor excitation). This is due to the fact that acceptor emission spectral bleedthrough in the donor channel is generally negligible, provided the donor emission filter is properly chosen. As a benefit, the necessary calibration of the system (*i. e*., IRF) or background correction are much simpler to achieve in these conditions.

### 2.5 Double-stranded DNA sample preparation

NIR dyes were obtained from ThermoFisher Scientific (Waltham, MA). Oligodeoxynucleotides (oligo-DNAs) were synthesized and labeled by IBA Lifesciences (Göttingen, Germany). The sequences of two complementary oligo-DNAs were as in Ref. (53), with the “top” strand’s sequence given by 5’-TAA ATC TAA AGT AAC ATA AGG TAA CAT AAC GGT AAG TCC A-3’. Alexa Fluor 700 (AF700) and Alexa Fluor 750 (AF750) were used as NIR FRET pair. The donor (AF700) was conjugated to dT at position 1 of the top strand, and the acceptor (AF750) was conjugated to dT at three separate positions (17, 22 and 27) on the “bottom” strand for each FRET sample respectively. All purchased fluorescently conjugated oligo-DNAs were provided purified using high-performance liquid chromatography (HPLC) and lyophilized. Unlabeled strands were purified using desalting method and delivered as lyophilized form. Each lyophilized oligo-DNA was first resuspended with Tris-EDTA buffer pH 8 (Sigma-Aldrich, MO) to make a 100 nM stock solution. To perform the hybridization, AF700 oligo-DNA strands were mixed with AF750 oligo-DNA strands at 1:1 molar ratio at 50 nM final concentration for 100 μL reaction volume. For donor-only and acceptor-only samples, unlabeled oligo-DNAs were used as complementary strands at 1:1 molar ratio. The mixture of oligo-DNAs was heated at 95 °C for 5 min using dry heating block and cooled at room temperature for 30 min to obtain a mixture of double-stranded DNS (dsDNA) and residual unhybridized single-labeled oligo-DNAs. Identical samples were used for the IVIS and the wide-field MFLI measurements.

### 2.6 Animal experiments

All animal procedures were conducted with the approval of the Institutional Animal Care and Use Committee at both Albany Medical College and Rensselaer Polytechnic Institute. Animal facilities of both institutions have been accredited by the American Association for Accreditation for Laboratory Animals Care International. Athymic female nude mice were purchased from Charles River (Wilmington, MA). All animals were in healthy condition. Tf probes were prepared by conjugating iron-bound Tf with fluorophores per manufacturer’s instruction. AF700 and AF750 were used as donor and acceptor, respectively. The animals were anesthetized with isoflurane before being retro-orbitally injected with 40 μg of AF700-Tf (donor) and/or 80 μg of AF750-Tf (acceptor) conjugates and imaged immediately. The intensity-based measurement lasted approximately 2 hours (2 s per channel, ≈ 34 seconds between time-point). The time-resolved data were acquired continuously for 90 minutes (≈ 43 seconds per acquisition). Each intensity FRET measurement involved three mice. The single-donor mouse was injected with AF700-Tf, the single-acceptor mouse with AF750-Tf and the double-labeled FRET mouse was injected with a mixture of AF700-Tf and AF750-Tf at acceptor:donor (A:D) ratio of 2:1 (40 μg of AF700-Tf and 80 μg of AF750-Tf). The lifetime measurement used only one mouse injected with a mixture of donor and acceptor. During imaging, mice were kept anesthetized using isoflurane, and their body temperature maintained using a warming pad (Rodent Warmer X2, Stoelting, IL) on the imaging plane.

### 2.7 Immunohistochemistry

Mice were injected with 40 μg Tf-biotin conjugates (Sigma-Aldrich, Inc., MO) or PBS buffer and sacrificed 6 hr post-injection. Bladders were collected, fixed in 4 % paraformaldehyde for 24 hr and processed for embedding and sectioning (43). Tissue sections were analyzed by immunohistochemistry using ABC Elite and NovaRed peroxidase substrate kit (Vector laboratories, CA) to visualize Tf-biotin. Parallel bladder sections were stained with Hematoxylin and Eosin and imaged using a 10x magnification microscope for tissue morphology visualization.

### 2.8 FRET quantification using decay fits of MFLI data

#### 2.8.1 dsDNA samples

*f*_*D*_ *E* (product of the fraction of donor involved in FRET and the FRET efficiency of the FRET sample) was quantified by fitting the fluorescence decays in each pixel of selected regions of interests (ROIs) to a bi-exponential model (Eq. 8) and retrieving the amplitude-weighted averaged lifetime ⟨τ⟩_*a*_ (Eq. 10). IRFs were acquired using a sheet of white paper as sample after removing all emission filters. After convolution, the tail portion of each pixel’s decay (99%-2% of the peak value) was fit using the MATLAB function *fmincon*() for least squares minimization of the cost function or, alternatively, the non-linear Levenberg-Marquardt algorithm implemented in the free software AlliGator (46, 61). After ⟨τ⟩_*a*_ was calculated for every decay of interest (including donor-only and double-labeled FRET sample), Eq. (12) was used to calculate *f*_*D*_ *E*.

#### 2.8.2 Dynamic Tf-TfR FRET *in vivo* imaging

The liver and bladder ROIs were delineated via intensity thresholding of the last time point in the series. Since the mouse did not move laterally along the imaging plane during the ≈ 90 min of imaging, the same mask was appropriate for all time-points. The donor-only lifetime was retrieved using the averaged mean-lifetime values ⟨τ⟩_*a*_ (Eq. 10) from the urinary bladder over the first five acquisitions (τ_0,*UB*_ = 1.03 ns). This method neglects environment-dependent changes of AF700-Tf lifetime between urinary bladder and liver, which measurements on AF700-Tf only injected mice discussed in Supplemental Note S1 indicate are minimal in this system. Note that this methodology might not hold in other situations (*e.g*., due to putative pH dependence of lifetime (51)) and a separate measurement with a donor-only labeled mouse may be needed to obtain the local donor lifetime (similar to the measurement described in Supplemental Note S1).

As discussed in Supplemental Note S1, we also verified that autofluorescence signal from tissues was negligible in the conditions of our experiments, and did not influence the analysis. We emphasize that this depends on the extrinsic signal intensity being significantly larger than the autofluorescence signal, and needs to be assessed for each experimental system as well as setup settings.

All other analysis steps and calculation of FRET efficiency were performed similarly as described above for the dsDNA sample, with the exception of the constraints for the two lifetimes τ_0_ and τ_1_, which were set to [0.2, 0.4] and [0.9, 1.1] respectively.

### 2.9 FRET quantification using sensitized emission analysis of IVIS data

#### 2.9.1 dsDNA samples

Background subtraction was performed on all excitation/emission channels. The correction factors (*d*_*D*_, *l*_*D*_, *d* _*A*_ and *l* _*A*_) were then determined using Eq. A.12-A.13. Additionally, the *γ* correction factor was determined using the known quantum yields and fluorescence emission spectra (Fig. S1) for the NIR dyes, as well as filter specifications and camera quantum efficiency of the IVIS imaging setup. The calculated value was 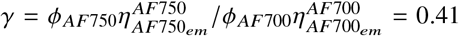. Afterwards, *f*_*D*_*E* was calculated according to Eq. 7.

#### 2.9.2 Dynamic Tf-TfR FRET *in vivo* imaging

The correction factors (*d*_*D*_, *l*_*D*_, *d* _*A*_ and *l* _*A*_) were determined in a dynamic fashion at each time-point using the intensities of the liver and bladder ROIs of all three mice and Eqs. A.12-A.13 (Fig. 2). Intensity-based *f*_*D*_ *E* was then calculated as described above for the dsDNAs.

## 3 RESULTS

### 3.1 Short double-labeled dsDNA strands as FRET standards

Double-labeled double-stranded DNA (dsDNA) molecules provide a simple and convenient way to design molecules with well-defined distances between donor and acceptor fluorophores that can be used as FRET standards (62, 63). In an ideal case, the base pair separation between donor and acceptor dyes determines the FRET efficiency based on the B-DNA model structure. The larger the separation, the lower the FRET efficiency, which depends on the ratio of the distance between the two fluorescent donor and acceptor fluorophore molecules to to their Förster radius *R*_0_ (*R*_*AF*700/*AF*750_ = 7.8 nm). For this study, three dsDNA FRET standard samples were prepared by hybridization of donor-or acceptor-labeled complementary 35 oligonucleotide long single-stranded DNA (ssDNA) molecules characterized by donor-acceptor distances of 17, 22 and 27 nucleotides (Fig. 1A) (53). The fluorophores used here (Alexa Fluor 700 and Alexa Fluor 750) are near-infrared (NIR) emitting fluorophores widely adopted for *in vivo* imaging applications (41, 64). The donor fluorophore (AF700) is located at the end of the same ssDNA, while the acceptor (AF750) is located in different positions of the complementary strand, but surrounded by a common nucleotide pattern, in order to ensure a constant environment (and therefore a common Förster radius) for all samples (53). These dsDNA samples were imaged for both FLI- and intensity-based FRET analysis using MFLI and IVIS imagers, respectively.

**Figure 1:**
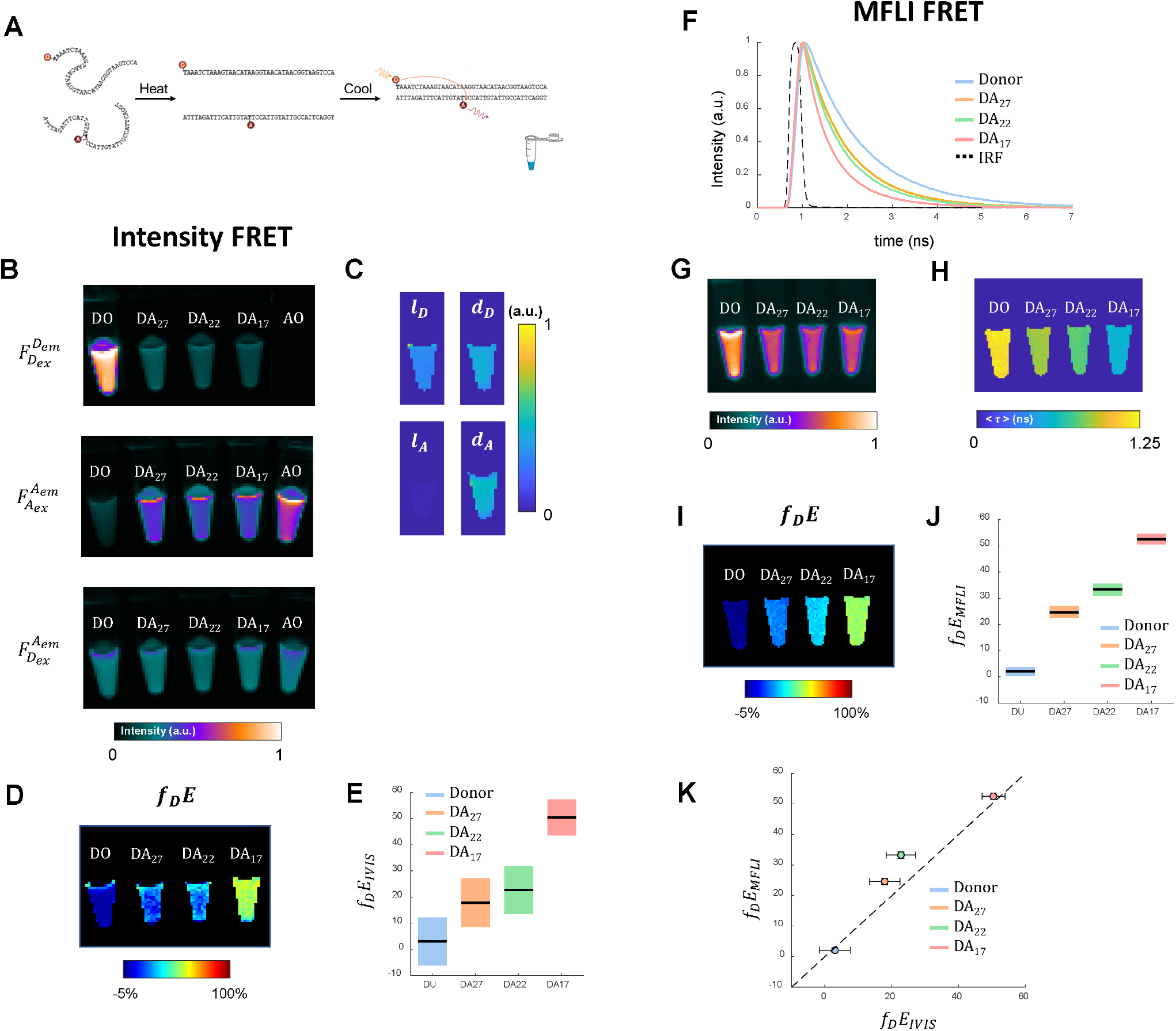
*in vitro* comparison of intensity- and MFLI-FRET imaging methods. A: Oligonucleotide sequences used for hybridized DNA FRET sample *D A*_17_. The donor, Alexa Fluor 700 was conjugated to dT at position 1. The acceptor, Alexa Fluor 750, was conjugated to dT positions 17, 22 and 27 on the complementary strand. Hence, the distance between the donor and the acceptor for “DA 17” after hybridization was 17 base pairs, which corresponds to approximately 5.8 nm. L, Scatter plot of *f*_*D*_ *E* results (mean ± standard deviation) retrieved through intensity- and FLI-FRET. B: Fluorescence intensity data acquired with the IVIS Lumina XRMS Imaging setup: donor only (DO) dsDNA, acceptor only (AO) and FRET samples (*D A*_17_, *D A*_22_ and *D A*_27_) imaged with donor channel 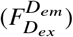, acceptor channel 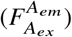 and FRET channel 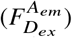. Results from DNA FRET standard sample. C: Spectral correction coefficient maps. D: *f*_*D*_ *E* map retrieved through intensity FRET. E: Boxplot of *f*_*D*_ *E* values retrieved using intensity FRET. F: Normalized MFLI decays measured from the donor-only and FRET dsDNA samples (whole vial ROI). G: Max-normalized intensity measurements using a gated-ICCD. H: Amplitude-weighted mean lifetime of the donor-only and FRET dsDNA samples. I: *f*_*D*_ *E* map retrieved through lifetime-based FRET. J: Boxplot of *f*_*D*_ *E* values retrieved using lifetime-based FRET. K: Comparison of intensity-(horizontal axis) and lifetime-(vertical axis) based mean FRET values.

**Figure 2:**
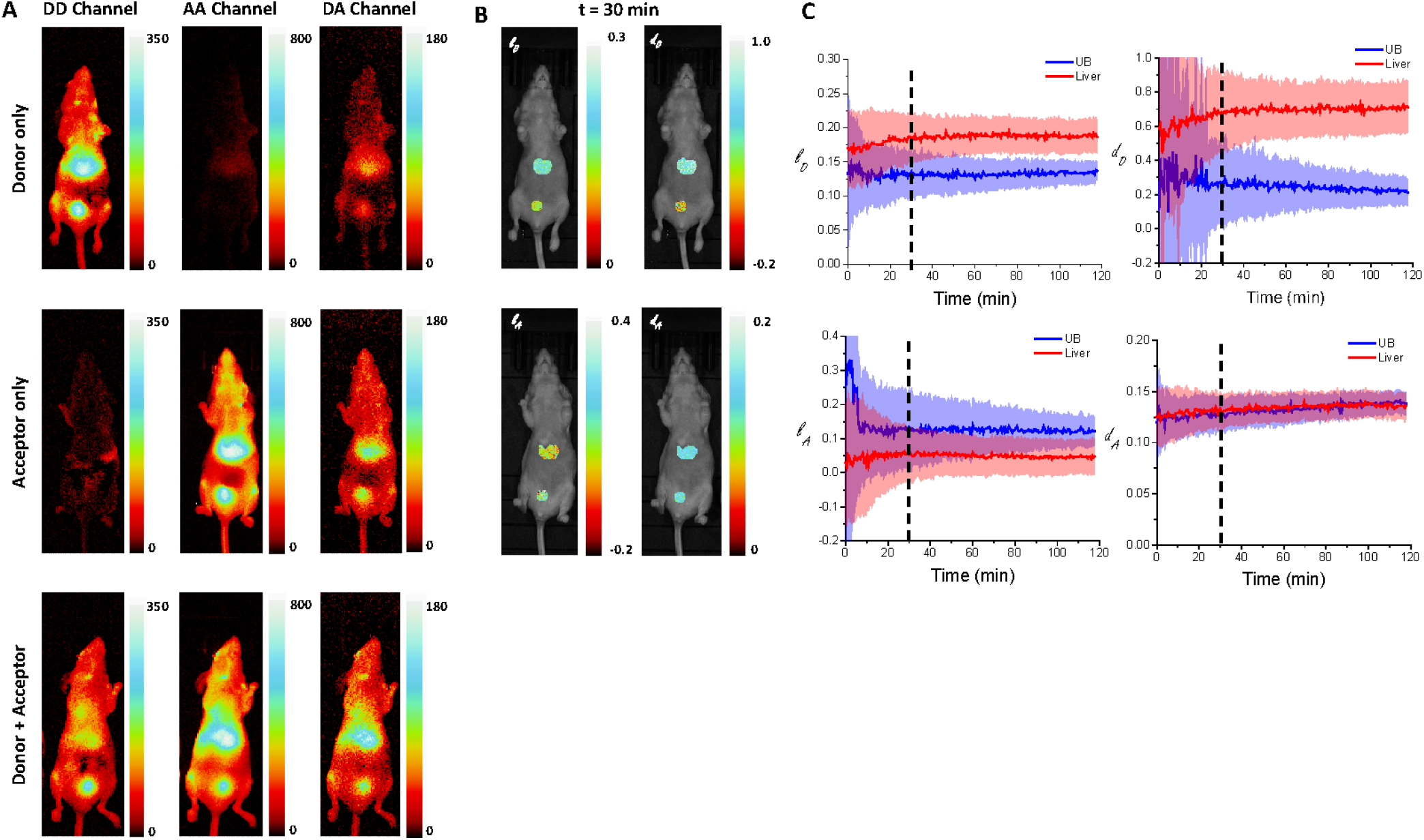
Dynamic SE-FRET spectral correction *in vivo* using IVIS. A: 3-cube (background-corrected) image data of mice injected with NIR-Tf, 30 minutes post injection (p.i.). Top row: donor-only mouse, middle row: acceptor-only mouse, bottom row: donor + acceptor mouse (acceptor-to-donor ratio = 2:1). Each row is comprised of a 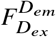 (DD Channel), 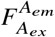 (AA Channel) and a 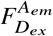 (DA Channel) image, as indicated on the top of each image column. The intensity is color-coded as indicated by the color bar shown in the top right of each column A. B: Spectral correction coefficient maps for t = 30 min *p.i*. for the liver and bladder ROIs as indicated in the top right corner of each map. The *l*_*D*_ and *d*_*D*_ maps are computed from the images shown in the top row (donor-only mouse), while the *l* _*A*_ and *d* _*A*_ maps are computed from the images shown in the middle row (acceptor-only mouse), color-coded according to the color bar shown on the right of each map. C: Temporal evolution of the spectral correction coefficients in both ROIs. The plots show the average and standard deviation of each coefficient in the liver (red curve and red shaded area) and the bladder (blue curve and blue shaded area), corresponding to the respective maps shown in B. The 30 min *p.i*. time point illustrated in panels A & B is indicated by a dashed vertical line in all plots.

In contrast with MFLI, which requires the imaging of one or two samples only (FRET, *i.e*. donor + acceptor and optionally, donor-only), three samples are necessary for intensity FRET analysis: donor-only, acceptor-only and FRET sample (referred to as donor-acceptor *n*, or, “*D A*_*n*_”, where *n* indicates the number of nucleotides separating donor and acceptor). All samples were imaged with donor, acceptor and FRET excitation/emission channels, with the same field of view and excitation power settings (Table 1). The fluorescence intensity maps of all samples from all channels are shown in Fig. 1B. Correction factors *d* _*A*_ and *l* _*A*_ were obtained from the acceptor-only dsDNA sample, while corrections factor *d*_*D*_ and *l*_*D*_ were obtained from the donor-only dsDNA sample (Fig. 1C, Table S3). The correction factor *γ*, was calculated using the known quantum yield of both dyes, as well as by calculating detection efficiency with known emission spectra and detection wavelength bands used herein 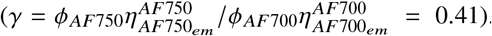. Using this information, calculation of the FRET efficiency of the sample using Eq. 7 resulted in *f*_*D*_ *E* values shown in Fig. 1C, D (Table S4).

The fluorescence decays and the intensity, average lifetime, and *f*_*D*_ *E* maps of all samples from all channels are shown in Fig. 1F-I. The amplitude-weighted mean lifetime of a dsDNA donor-only sample consisting of the donor-labeled ssDNA hybridized to an unlabeled complementary ssDNA strand was measured as τ_*DO*_ = 1.19 ± 0.05 ns. Using the amplitude-weighted mean lifetimes of the double-labeled dsDNA samples and Eq. 12, led to the values of *f*_*D*_ *E* represented in Fig. 1I. Fig. 1K compares the MFLI results to those obtained with the IVIS system. We hypothesize that the difference between both sets of results is due to residual donor bleedthrough when excited with the acceptor wavelength (see Fig. 1B, 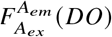. To properly correct for this, the measurement of 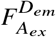(X = DO, AO and the 3 FRET samples) would be needed and additional correction factors included in the analysis. This further highlights the complexity of the intensity-based FRET approach for these applications.

The larger standard deviation of the intensity-based FRET results compared to the lifetime-based results noticeable in Fig. 1L is likely due to the lower SNR of the IVIS data. Another possible contributor is the fact that average correction factors were used, as they were computed using different control tubes (DO and AO), located at different positions and angles than the *D A*_*n*_ tubes.

Importantly, these complications are not present for FLI-FRET quantification, and we observed low standard deviation across the vial ROIs (see Fig. 1I, J).

### 3.2 *in vivo* FRET imaging of transferrin-transferrin receptor binding

We next demonstrated dynamic monitoring of ligand-receptor engagement *in vivo* using sensitized emission FRET and compare it to MFLI-FRET.

The Tf-TfR system was chosen as a model for *in vivo* FRET imaging of ligand-receptor engagement since transferrin has been used widely as a carrier for drug delivery (65). Tf-TfR binding was monitored by either IVIS imaging according to intensity FRET imaging protocol as described in Material & Methods or using the MFLI imager as described previously (42). Briefly, the animals were intravenously injected with NIR-Tf fluorescently labeled probes and imaged continuously for over an hour and a half at 30 to 43 sec interval steps depending on the instrument. As previously observed, there was a significant increase in fluorescence accumulation in the liver and the urinary bladder as a function of time, while very little was detected in other organs (43, 44, 64). This finding was consistent across the intensity and the lifetime FRET measurements.

#### 3.2.1 Sensitized emission FRET quantification using IVIS data

As shown in Fig. 2A & 3A, single-labeled donor-only or acceptor-only mice showed fluorescence intensity in the urinary bladder and the liver in *all* channels. As expected, donor-only mouse fluorescence was negligible in the acceptor channel, and acceptor-only mouse fluorescence was negligible in the donor channel. However, fluorescence intensity levels are clearly detected in the FRET channel (donor excitation and acceptor detection) for all organs of the donor-only and acceptor-only single-labeled mice, indicating significant spectral bleedthrough.

**Figure 3:**
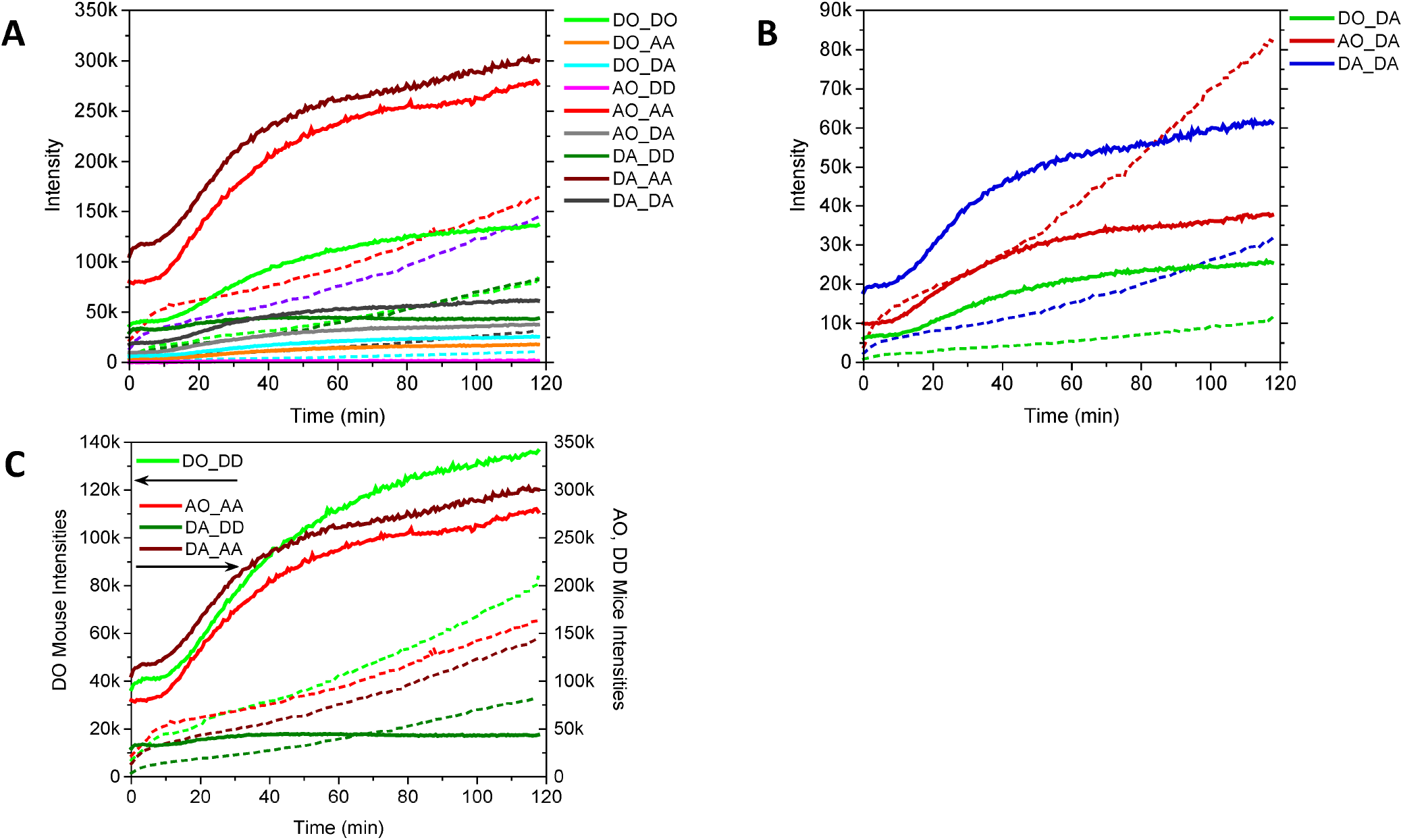
IVIS intensities observed in the different organs of the different mice (DO, AO & DA) used in the experiment. A: All measured intensities as a function of time. The graph legend refers to the different plots as *mouse_channel* (UB: thin dashed curves, liver: thick curves). B: FRET channel intensities variations observed in the urinary bladder (thin dashed curves) and liver (thick curves) of the 3 mice. The fact that DO_DA (FRET signal in the donor-only mouse) is non zero indicates donor bleedthrough in the acceptor detection channel, while the presence of AO_DA signal (FRET signal in the acceptor-only mouse) indicates direct excitation of the acceptor with the donor excitation wavelength. The signals in the UB of all mice behave qualitatively similarly, as are their liver signals, demonstrating that DA intensity only is not a sufficient signature of FRET. C: Pharmaco-kinetics of the probes *not* affected by FRET in each organs are similar in the 3 animals (referred to as DO_XX, AO_XX and DA_XX, where is XX designate the excitation/detection channel). In the liver (thick curves), the donor probe in the DO mouse (DO_DD, light green), the acceptor probe in the AO mouse (AO_AA, red) and the acceptor probe in the FRET mouse (DA_AA, wine) all show a lag phase followed by a rapid rise and a plateauing of the observed intensity. Similarly, in the urinary bladder, these probes (same colors, thin dashed curves) show a first rapid increase followed by a slower accumulation. By contrast, and as expected, in the FRET mouse, the donor signal observed in the liver (DA_DD, thick dark green curve) behaves very differently, with almost no lag phase and a slow decrease throughout most of the observation. Meanwhile, the same donor signal in the bladder (DA_DD, thin dashed dark green curve) follows a similar trend as the other probes in the other mice, *a priori* indicating no particular interaction going on in this organ. Note that this graph has two intensity axes, the left one used for the D-only mouse and the right one for the two others, as indicated by the arrows, due to different ranges involved.

In the double-labeled mouse, all channels, including the FRET channel 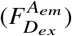, showed fluorescence intensity in the liver as well as in the urinary bladder (bottom row of Fig. 2A). Both acceptor and FRET channel intensity measurements showed an accumulation of fluorescence probe in the liver and the urinary bladder over time, which was qualitatively similar to that observed in the single-labeled mice (Fig. 3B & C). The only striking difference with these “control” mice was observed in the donor channel (Fig. 3C), where the bladder signal increased continuously, as in the other mice, but the liver signal rapidly plateaued and eventually started decreasing.

These results are qualitatively consistent with FRET occurring upon ligand-receptor interaction in the liver, as expected based on its role in iron metabolism and the known high level of TfR expression in the liver. By contrast, the common increasing FRET channel intensity in the urinary bladder of all mice suggests that this signal was mostly contributed by donor spectral leakage and/or direct excitation of the acceptor. However, the only way to exclude any possibility that FRET was occurring the urinary bladder of the FRET mouse, was by performing sensitized emission FRET analysis in a dynamic fashion to account for spectral bleedthrough at each time-point.

Herein, we used the data from the single-labeled (DO and AO) mice for spectral correction of the FRET mouse data (*i.e*., mouse injected with a mixture of donor and acceptor-labeled Tf). As noted above, the pharmaco-kinetics of the probes appeared very similar in both organs for all 3 mice (Fig. 3C), supporting the use of information from the donor-only and acceptor-only mice to obtain the four correction coefficients for acceptor and donor spectral crosstalk and bleedthrough for the donor-acceptor mouse analysis (Fig. 2B). All coefficients were calculated and applied at each time-point (Fig. 2C).

It is noteworthy that all coefficients are tissue-dependent and vary over time. The tissue dependency is not unexpected based on their definitions A.12 & A.13, which involves either excitation intensities or detection efficiencies. Both might be dependent on tissue depth and the optical properties of intermediate tissues. Since these properties are wavelength-dependent, the temporal variation of ratios of these coefficients likely reflects the changing depth distribution of the probes as a function of time.

In addition, we applied the same *γ* correction factor as used for the dsDNA FRET samples analysis. Because this constant factor does not fully correct for the wavelength dependence of optical absorption variation in biological tissue (and the changes in fluorophore distribution as a function of time, as discussed above), this assumption might contribute to additional uncertainties in the intensity corrections.

Using these parameters and the intensity in the FRET channel, *f*_*D*_ *E* was calculated at each time point in the liver and urinary bladder of the donor-acceptor mouse. Intensity-based dynamic *f*_*D*_ *E* of Tf-TfR ligand-receptor interaction showed increasing mean FRET efficiency in the liver and no significant *f*_*D*_ *E* in the urinary bladder (*f*_*D*_ *E* = 1.6 ± 1.6%, Supplemental Fig. S3A) throughout the 2 hr duration of the observation (Fig. 5B). This result directly correlates with the known physiology of Tf, which binds to its receptor in the liver allowing FRET to occur. The absence of FRET in the urinary bladder indicates excretion and inability to bind TfR, of degraded Tf or their degradation products (free fluorophores). It is worth noting that *f*_*D*_ *E* values retrieved over the first 20 minutes in the mouse liver are negative (Fig. 5B). This is due to negative *F*^*FRET*^ values calculated according to Eq. 4 during this time-interval, and most likely indicates inadequate correction factors. While the pharmaco-kinetics of the two probes (AF700-Tf and AF750-Tf) in the two “control” mice are similar to those in the FRET mouse (Fig. 3C), they are not identical, reflecting unavoidable inter-animal physiological variation. Moreover, due to potential difference in probe distribution and tissue properties in the different mice, discrepancies between correction factors computed in one mouse and used in another are further expected to contribute to these erroneous results.

#### 3.2.2 Lifetime-based FRET quantification using MFLI data

In comparison, lifetime-based FRET analysis was much more straightforward. As discussed next, MFLI data from the donor emission only (695 nm excitation, 721 ± 6 nm detection) from a double-labeled mouse was sufficient to evaluate ligand-receptor interaction.

Similarly to the experiment involving intensity measurements (Fig. 4A, blue curve), a steady increase of donor fluorescence was observed in the urinary bladder (Fig. 4B, blue curve). Intriguingly, contrary to the case of the intensity-based experiment, where the donor channel liver fluorescence plateaued and slightly decreased toward the end of the measurement (Fig. 4A, red curve), the MFLI donor channel intensity in the liver steadily decreased over the span of the measurement Fig. 4B, red curve).

**Figure 4:**
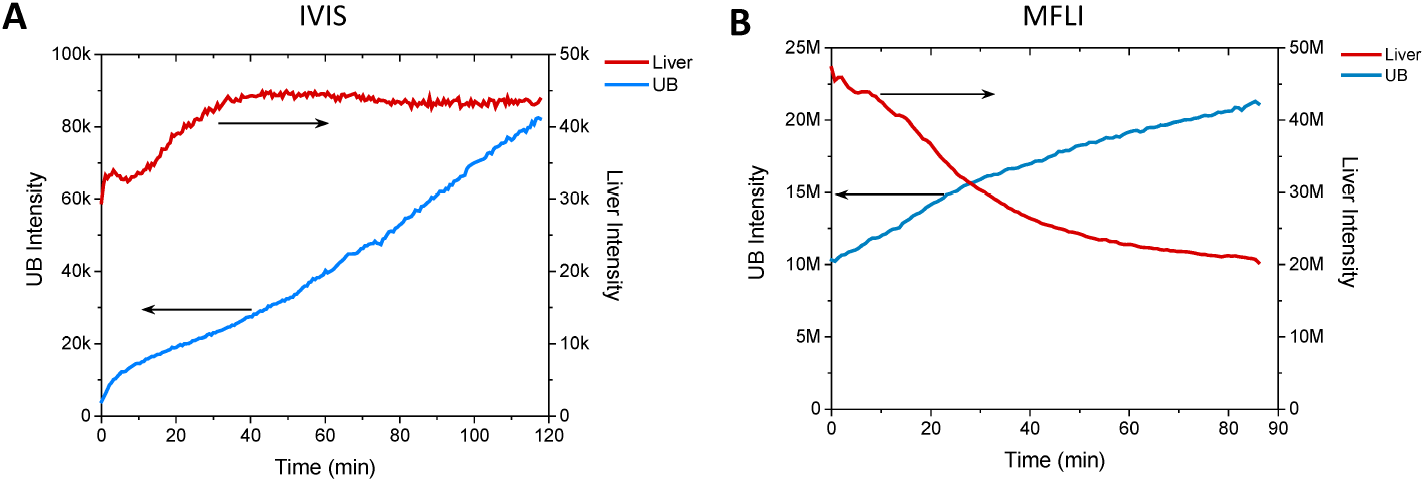
Comparison of donor channel intensity dynamics: IVIS vs MFLI. A: Donor channel intensity plots for the liver (red) and urinary bladder (blue) ROIs of the double-labeled mouse observed with the IVIS system. B: Donor channel integrated intensity plots for the liver (red) and bladder (blue) ROIs of the double-labeled mouse observed with the MFLI system. Both datasets exhibit some decrease of the liver donor channel signal after some time, which further analysis described in the text links to increasing FRET in the liver.

**Figure 5:**
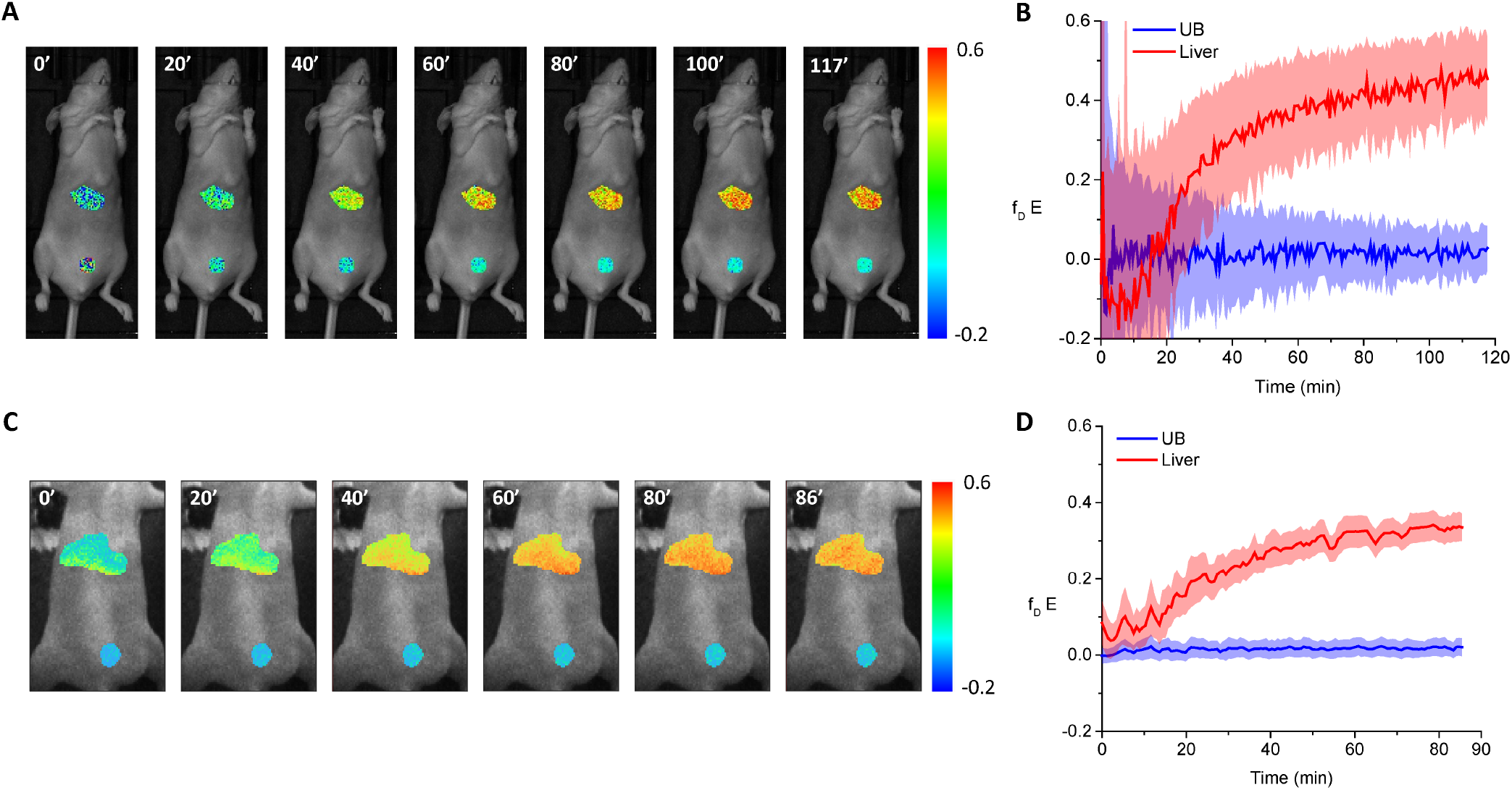
*In vivo* comparison of intensity- and MFLI-FRET dynamic imaging. A: Maps at selected time points of intensity-based *f*_*D*_ *E* in the liver and the urinary bladder of a mouse injected with Tf A:D 80:40 μg (total time points: 210). B: Corresponding intensity-based *f*_*D*_ *E* time trace. C: Maps at selected time points of lifetime-based *f*_*D*_ *E* in the liver and the urinary bladder of a mouse injected with Tf A:D 80:40 μg (total time points: 127). D: Corresponding lifetime-based *f*_*D*_ *E* time trace. For B and D, solid lines mark the average value and the shaded areas indicate the standard deviation across all pixels within each organ ROI.

Because no acceptor or FRET channel information is available in the MFLI measurement, we proceeded with the donor lifetime analysis described in Material and Methods and illustrated previously with the DNA sample. This analysis requires a donor-only lifetime to which to compare the mean donor lifetime observed in the region of interest. As discussed in Material and Methods and Supplemental Note S1, the mean donor lifetime measured in the urinary bladder during the first few time points of the experiment (τ_*bladder*_ = 1.03 ns) was used for both urinary bladder and liver ROI throughout the experiment. This ignores potential differences between donor lifetime in the two environments and any possible changes of these lifetimes across time. Control experiments performed in donor-only labeled mice indicate that this assumption is reasonable (Supplemental Note S1 and Supplemental Fig. S6). Moreover, as discussed in Supplemental Note S1 and Supplemental Fig. S6, a separate analysis performed using a time-dependent donor-lifetime resulted in similar conclusion to the one obtained using this simpler assumption.

This analysis suggests that MFLI FRET measurements in other biomedical assays that do not provide a validated internal negative FRET control such as the urinary bladder in the present experiments, can instead use the long lifetime component obtained from bi-exponential fitting of tissues or organs in which FRET takes place, in so far as it matches that observed in similar experiments performed with a donor-only-labeled mouse, as indicated in Table S5.

Lifetime FRET analysis of the donor fluorescence decay curves with bi-exponential fitting yielded amplitude-weighted average lifetimes ⟨τ⟩_*a*_ from which *f*_*D*_ *E* values were calculated using Eq. 12. *f*_*D*_ *E* in the liver is increasing over time due to the combined high expression of TfR in that organ and the large vascular fenestration (100-175 nm) of hepatic sinusoids, facilitating passive accumulation of Tf in the liver (Fig. 5C-D) (66). By contrast, *f*_*D*_ *E* within the urinary bladder was negligible throughout the imaging session (*f*_*D*_ *E* = 1.7 ± 0.3% Supplemental Fig. S3B), consistent with the continuous increase of donor fluorescence intensity observed in that organ (Fig. 4B, blue curve). This finding suggests that there was negligible FRET in the urinary bladder (either low FRET efficiency *E*, or low FRET-ing donor fraction *f*_*D*_, or both). This observation indicates that either (i) there was no or negligible binding of the Tf present in the urinary bladder to TfR, or (ii) that the observed donor fluorescence was a result of degraded AF700-Tf, leading to free fluorophore accumulation. These hypotheses are both supported by immunohistochemical analysis of bladder tissues from mice intravenously injected either with biotin-labeled Tf or PBS (negative control, Fig. S2) showing no Tf staining in the lining of all bladders.

Overall, these results indicate that bladder tissues do not accumulate TfR-bound Tf upon intravenous injection. In any case, excluding the negative *f*_*D*_ *E* values retrieved over the first 20 minutes in the intensity FRET analysis for the liver, *f*_*D*_ *E* quantification obtained by both approaches are in good agreement with each other, considering that the measurements were performed using different mice (and in the case of the intensity-based FRET analysis, multiple mice were used to obtain the different correction parameters). Interestingly, quantitative analysis of the two kinetic curves shown in Fig. 5B-D using a simple exponential model (Supplemental Figure S4) yielded similar time constants τ_*kin*_ in both experiments (intensity-based analysis: τ_*kin*_ = 25.3 ± 0.8 min, lifetime-based analysis: τ_*kin*_ = 24.5 ± 1.1 min), consistent with those observed in similar experiments (46).

## 4 DISCUSSION

Noninvasive molecular imaging approaches have been used for assessment of drug distribution and delivery *in vivo* with great success (67, 68). Noninvasive imaging enables longitudinal assessment of preclinical drug candidates without the need to sacrifice animals at every time point of interest. Moreover, using the same animal across multiple time points minimizes inter-animal variation (Fig. 6A). The localization of imaging contrast agents provides insight into the distribution of pharmaceutical or biopharmaceutical compounds administered to the animals. Hence, molecular targeted imaging has been used for *in vivo* studies of pharmaco-kinetics and drug distribution using nuclear imaging (PET and SPECT) (69, 70). The output of these modalities is intensity information, which is used to represent the localization of the drug. Unfortunately, this information cannot be used to distinguish between co-localization in the same spatial region and the accurate direct measurement of target binding or cellular delivery. This limitation often requires an invasive *ex vivo* approach to fully reveal binding of the administered compound to its respective target. The method of choice is histopathology – including immunohistochemistry or immunofluorescence staining (Fig. 6A). Though, analysis of *ex vivo* samples lacks whole-body drug distribution context, which should include other important organs besides the pathological ones. Additionally, *ex vivo* investigations require sacrifice of the animal for each time point considered, leading to increased biological variations.

**Figure 6:**
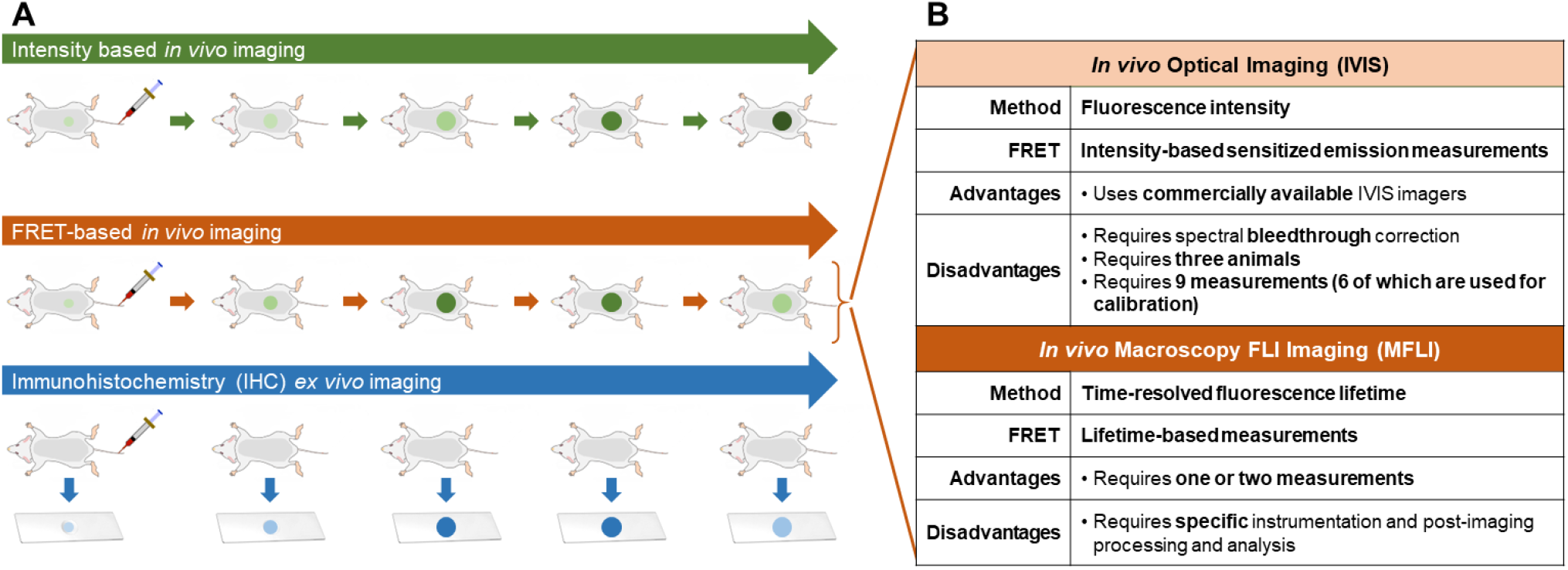
Comparison of different approaches for longitudinal preclinical molecular interaction monitoring. A: Conventional preclinical longitudinal studies such as IHC (bottom, blue color scheme) require invasive analysis at each time point, thus increasing the number of animals used and introducing inter-animal variability. In contrast, preclinical longitudinal molecular imaging (top, middle; green and red color schemes) enables the use of the same animals throughout the whole experiment, reducing the number of animals and minimizing inter-animal variability. Circles with different contrast indicate location of probe accumulation and recorded signal intensities. B: Intensity-based imaging cannot discriminate bound-vs. non-bound-probes, measuring only the passive accumulation of probes in the organ of interest. In contrast, FRET *in vivo* imaging can measure dynamic receptor-ligand engagement using either intensity-based sensitized emission or lifetime-based *in vivo* imaging approaches.

We have previously shown that dynamic TfR-Tf receptor-ligand engagement can be studied *in vivo* using MFLI FRET imaging (43, 44, 64). Transferrin (Tf) is an iron-carrying protein which can bind its homodimeric transferrin receptor (TfR) at the surface of all cells in the organism. TfR is a homodimeric membrane-bound glycoprotein characterized by an inter-dimer distance less than 10 nm, which allows the monitoring of Tf binding using FRET (64). Tf-TfR binding has been monitored both *in vitro* and *in vivo* using FLI FRET imaging and validated by immunohistochemistry (43, 44, 46). As NIR-Tf probes are introduced into the body via intravenous injection, they will primarily label the liver, which acts as a major location for iron homeostasis regulation and displays a high level of TfR expression. Free dye and/or small labeled degradation products of NIR-Tf probes end up accumulating non-specifically in the urinary bladder, due to its role as an excretion organ (46).

As demonstrated here, this type of study can be performed using both intensity- and lifetime-based approaches (Fig. 6). However, the intensity-based approach requires spectral correction that are cumbersome to implement experimentally, as additional calibrating samples are necessary. The intrinsic complexity of the corresponding 3-cube method commonly employed *in vitro* is further compounded by the fact that different animals need to be employed, raising questions about the reproducibility of this approach. Altogether, 9 independent measurements involving three mice were required (Table S5; Fig. 6B). While the data obtained with the two mice injected with donor-only and acceptor-only probes can in principle be reused for correction of new measurements with new mice injected with both probes, this requires that no change in acquisition parameters (and setup) takes place from one experiment to another, which might be difficult to ensure. In practice, it would be recommended to repeat these calibration measurements each time, increasing the cost and complexity of these measurements. Moreover, for the correction factors defined in Eqs. A.12 and A.13 to be valid, it is critical that the dye environment in the different mice used for their estimation, as well as the properties of the surrounding tissues, are similar (as implicitly assumed in the derivation). This may prove extremely difficult to ensure due to mouse-to-mouse variability, in particular when perturbations such as tumor xenografts are involved, since xenografts grown from the same cell line often possess variable size, cell density and microenvironments across different animals (43, 71).

In *vivo* FRET measurement protocols have traditionally relied on reporting a relative increase in acceptor intensity of FRETing sample compared to non-FRETing sample (*i.e*., proximity ratio). However, imaging throughout the body of small animals upon probe injection, results in significant variation of fluorescence intensity as well as confounding emission leakage. Therefore, as in microscopy, intensity-based FRET *in vivo* macroscopy imaging should use the sensitized emission method, in which confounding emission leakages are properly corrected. However, sensitized emission FRET approach has not been adopted in small animal optical imaging, probably due to its complexity. Nevertheless, intensity-based sensitized emission FRET approach can be applied using widely available small animal imaging instruments such as the IVIS platform, making this imaging methodology accessible to many researchers. Sensitized emission FRET *in vivo* small animal imaging would allow the visualization of spatial drug distribution in a dynamic manner, enabling the understanding of the cellular mechanisms under pathophysiological context and providing valuable information for precision pharmaco-kinetics.

Lifetime-based FRET quantification provides robust and quantitative measures of target receptor engagement *in vivo* in a direct and non-invasive fashion. Even in the case of unique tumor microenvironments, lifetime-based FRET can be analyzed in each mouse independently. Hence, in its macroscopic implementation, MFLI-FRET is uniquely positioned to extract information regarding protein-protein interaction across entire small animals with high sensitivity (Fig. 6B). Importantly, MFLI FRET has been expanded to measure antibody-target engagement using NIR-labeled Trastuzumab, an anti-HER2 clinically relevant antibody in HER2 breast tumor xenograft models (44, 51). Therefore, NIR MFLI FRET imaging is a quantitative and non-invasive tool for the optimization of targeted drug delivery systems based on receptor-ligand or antibody-target engagement in tumors *in vivo*. MFLI-FRET should find broad applicability in *in vivo* molecular imaging and could be extended to applications as diverse as image guided-surgery or optical tomography as well as other antibody-target systems, including other HER or immunotherapy receptors. Considering the recent development of next-generation time-resolved SPAD cameras, which are simpler to use and more affordable than the gated-ICCD camera technology used in this study and have recently been validated in MFLI-FRET imaging of tumor xenografts in mice models of human breast and ovarian cancer (51), MFLI-FRET appears uniquely well-positioned to impact the field of molecular imaging.

## 5 DATA AND SOFTWARE AVAILABILITY

All data and results are available on a public cloud repository (72) in order to ensure reproducibility. Data corresponding to the mouse measured with the MFLI approach (Fig. 4B & Fig. 5C-D) has already been used in ref. (46), although not analyzed at the single-pixel level, as done in this work. Software used for analysis includes MATLAB (The MathWorks, MA), OriginPro (OriginLab, MA) and AlliGator, a free standalone software available on Github (46, 61). IVIS FRET Analysis, a simple LabVIEW software used to perform sensitized-emission FRET analysis of the mouse IVIS data as described in this article is available as source code on Github (73).

## 6 ACKNOWLEDGMENTS

This work was funded by the National Institute of Health (R01CA207725, R01CA237267, R01CA250636) and in part by HFSP grant RGP0061/2019 and NIH grant R01GM130942. XM wants to thank Shimon Weiss and the Weiss lab for their support during this work.

## 7 AUTHORS CONTRIBUTION

NS performed MFLI experiments. AR conducted IVIS imaging. JTS & XM analyzed the data. XM derived the equations. XI and MB designed and supervised the work. JTS, XM & MB wrote the manuscript. All authors reviewed the final manuscript.

## A APPENDIX: DERIVATION OF INTENSITY FRET EQUATIONS USED IN THE TEXT

### A.1 Definitions and notations

We use the notations of Ref. (53) with minor modifications. In particular, we drop the ‘ex’ and ‘em’ indices in the quantities 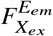 used in the main text, replacing them by 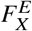 in order to simplify the notation.

As a reminder, data acquisition involves one of two types of excitation channels *X* (laser line in Ref. (53)), bandpass filter in the IVIS device), corresponding to the donor (*X* = *D*) or acceptor (*X* = *A*) excitation wavelengths, and two emission channels *E*, characterized by bandpass filters specific for the donor emission (*E* = *D*) or acceptor emission (*E* = *A*). 4 possible combinations of excitation and emission channels are therefore possible in principle: (*X, E*) ∈ {(*D, D*), (*D, A*), (*A, A*), (*A, D*)}. The last combination is rarely used in practice, as it is uncommon to observe emission in a wavelength band (*D*) shorter than the excitation band (*A*). While it could have been relevant to use it in the measurements described in the main text, no such data was collected, and therefore the formalism described next will ignore it.

For a given molecular species *S*, the signal collected using an (*X, E*) excitation/emission pair is denoted 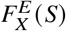, which we will assume to be corrected for background (data acquired in the same sample in the same conditions but with excitation source blocked).

Three different molecular species *S* are relevant in this study: *S* ∈ {*DO, AO, D A*}, where *DO* designates a donor-only species (molecule labeled with a donor fluorophore only), *AO* designates an acceptor-only species (molecule labeled with an acceptor fluorophore only) and *DA* designates a double-labeled (donor and acceptor) molecule. In principle, the *DA* species could be comprised of different sub-categories {*D A*_*i*_}_*i*=1…*n*_, characterized by different stoichiometries and/or different attachment sites and/or D-A distances. Formally, the same could be true of the *DO* and *AO* species, as fluorophore quantum yield could depend on the attachment site. In that case, we need to consider {*DO*_*i*_}_*i*=1…*d*_ and { *AO*_*i*_}_*i*=1…*a*_. We will here assume a single configuration for each species, but consider the case of different *DA* species in the last section.

Following previous notations (53), we further distinguish the physical process *Z* at the origin of the recorded signal using the notation 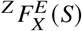. There are 3 processes of interest in this type of experiment:

i. *Z* = *D*: direct excitation of the donor, followed by donor emission,
ii. *Z* = *A*: direct excitation of the acceptor, followed by acceptor emission,
iii. *Z* = *D*→*A*: direct excitation of the donor, followed by non-radiative transfer to the acceptor, and acceptor emission.

We therefore have the following identities:

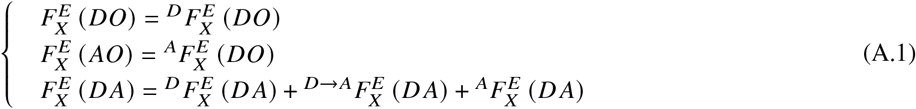

The first two simply state that no matter what excitation channel and emission channel are considered, only one single process needs to be considered when a single fluorophore species is present. The last identity expresses the fact that, in the case of the double-labeled species, three types of process can contribute to the signal: direct donor excitation/emission, donor excitation followed by FRET and acceptor emission, or direct acceptor excitation/emission.

Finally, for a sample comprised of a mixture of *N*_*D*_ donor molecules and *N*_*A*_ acceptor molecules, a fraction *f*_*D*_ (resp. *f*_*A*_) of which are part of a D-A pair, we have for the total number *N* of fluorophores in the sample, and the respective numbers *N*_*S*_ of species *S*:

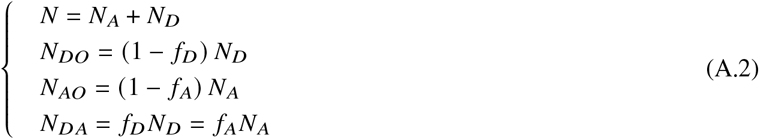

### A.2 Fundamental equations

The equations used in ref. (53) were defined for single-molecules and thus require a simple multiplication by one of the *N*_*S*_ factors and reintroducing the terms neglected in that work:

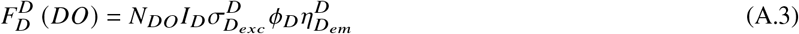

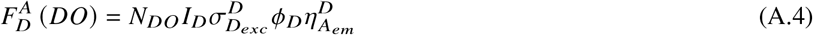

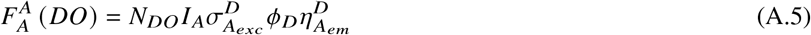

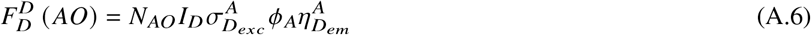

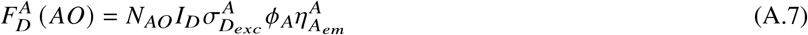

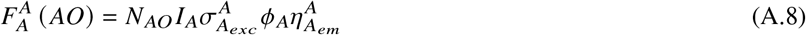

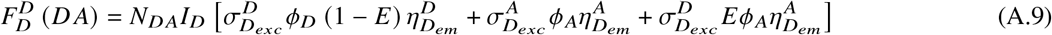

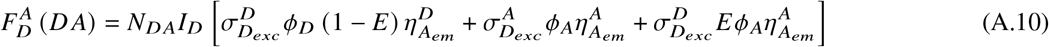

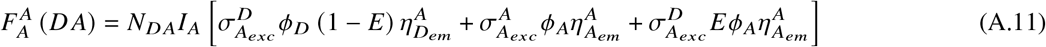

where Eqs. A.5-A.6 were assumed to be equal to zero in ref. (53) as were the last two terms of Eq. A.9 and the first and last term of Eq. A.11.

In the above equations:

- *I*_*X*_ is the *X*-excitation intensity (expressed in events per unit area, as detectors such as cameras do not measure photon energy, and instead only count the number of photon absorption events), which factors in integration time;
- 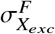 is the absorption cross-section of fluorophore *F* (= *D* or *A*) at wavelength *X*_*exc*_ (or the average absorption
- cross-section in the *X* excitation wavelength band);
- *ϕ*_*F*_ is the quantum yield of fluorophore *F* (= *D* or *A*);
- 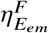 is the detection efficiency of fluorophore *F* (= *D* or *A*) in emission channel *E* (= *D* or *A*);
- *E* is the FRET efficiency of the *DA* pair.

Note that we ignore all 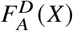 terms in this analysis (species excited with the acceptor wavelength and detected in the donor emission channel), as their contribution should be negligible in the present case, but some experimental situations might require their consideration to obtain fully corrected quantities.

Finally, these expression neglect any higher order photophysical effects such as re-excitation, saturation, etc., which could potentially play a role in some specific experimental situations but are deemed negligible here.

Based on these general equations, we can now look at the two “reference” samples measured in this study, namely donor-only (*DO, N* = *N*_*DO*_, *N*_*A*_ = *N*_*DA*_ = 0) and acceptor-only (*AO, N* = *N*_*AO*_, *N*_*D*_ = *N*_*DA*_ = 0). Using Eqs. A.3-A.5 we obtain:

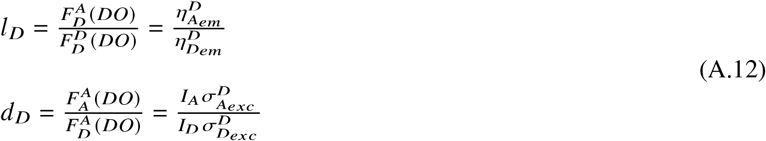

and using Eqs. A.6-A.8:

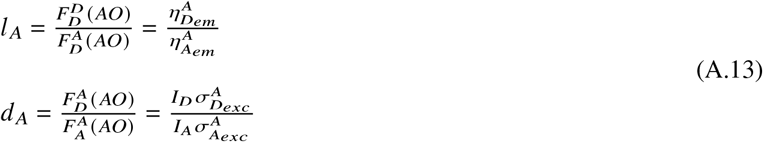

Coefficients *l*_*D*_ and *d* _*A*_ correspond to coefficients *l* and *d* in Ref. (53) and represent the donor leakage coefficient and the acceptor direct-excitation coefficient respectively. The two new coefficients *d*_*D*_ and *l* _*A*_ are counterparts of these coefficients, and are negligible if the donor absorption cross-section at the acceptor excitation wavelength, 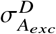, is negligible, and the detection efficiency of the acceptor in the donor emission channel, 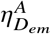, is negligible. These coefficients can be estimated from the *DO* and *AO* signals measured with the different excitation-emission (*X, E*) combinations, provided these quantities all correspond to the same integration time, and more generally, same detection parameters, as well as constant excitation intensity for a given (*X, E*).

### A.3 Pure *D A* sample case

The next step consists in extracting from Eqs. A.9-A.11, which are valid for “pure” *DA* species, an expression for *E* in terms of the measurable quantities 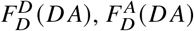 and 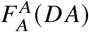. In order to simplify notations, we will define *D, A* and *F* as:

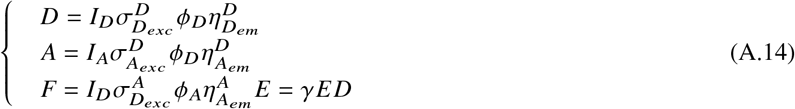

where the *γ* factor is defined by (53):

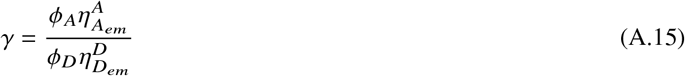

With these notations, Eqs. A.9-A.11 can be rewritten:

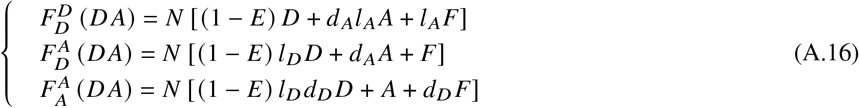

Replacing *F* by *γE D* results in 3 equations for the 3 unknowns *D, A* and *E*, the latter one being the only one of interest. Simple algebra yields:

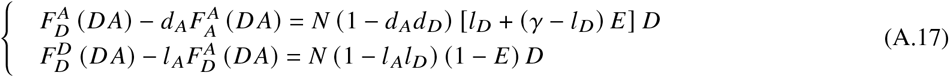

Taking the ratio of these two expression eliminates *N* and *D*, yielding the following result:

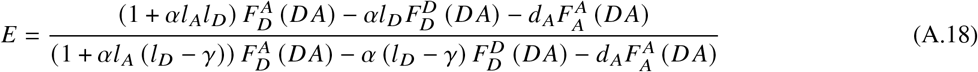

where we have introduced *α* defined by:

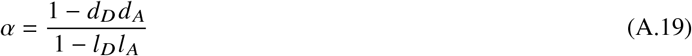

This formula is identical to Eqs. (10)-(11) of Ref. (53) when *d*_*D*_ = *l* _*A*_ = 0 (*α* = 1).

Note that Eq. A.19 can be expressed in terms of the sensitized emission FRET term *F*^*FRET*^ = *N F* and some additional terms. To express *NF* as a function of 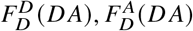 and 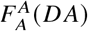, we look for (*u, v*) such that 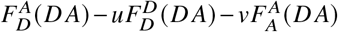 contains no *D* and *A* terms, based on the definitions of Eq. A.16. We obtain:

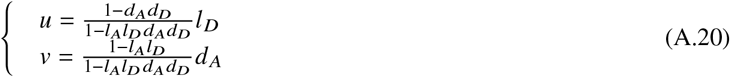

From this, we obtain the following expression for *F*^*FRET*^ = *N F*:

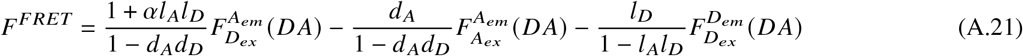

It is then straightforward to verify that *E* in Eq. A.18 can be rewritten:

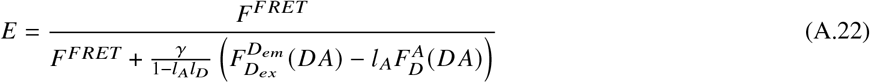

based on the definitions of Eq. A.16.

### A.4 Simple mixture case: D, A and DA mixture

When the sample is a mixture *M* as defined by Eq. A.2, the 3 measured signals are given by the sums:

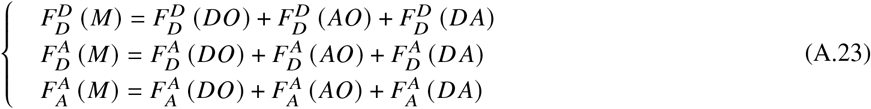

where the terms in the right hand side of Eq. A.23 are given by their expressions in Eqs. A.3-A.11. Using the definitions of Eq. A.14, we obtain:

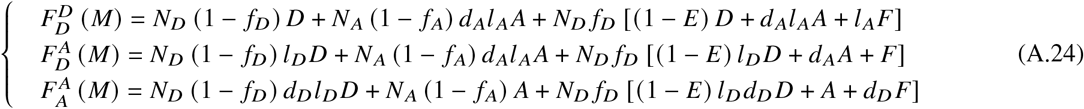

Using the identity *F* = *γE D* and the fact that *f*_*D*_ *N*_*D*_ = *f*_*A*_*N*_*A*_ (Eq. A.2), this can be rewritten:

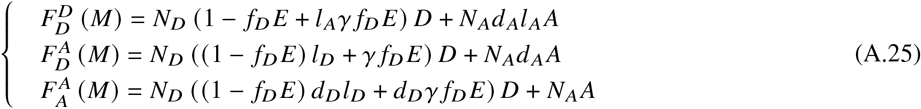

To eliminate *N*_*A*_*A*, the same combinations as in Eq. A.17 give:

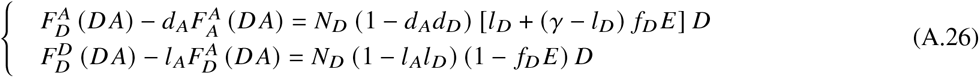

which are identical to Eq. A.17, except for the replacement of *N* by *N*_*D*_ and *E* by *f*_*D*_ *E*. The final result for the product *f*_*D*_ *E* is therefore identical to Eq. A.18, except for the quantities involved, which are now the intensities recorded for the mixture, rather than the pure *DA* sample:

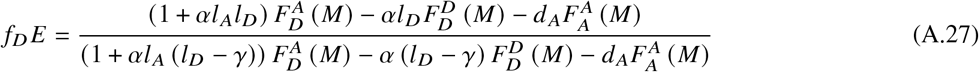

Note that this result is slightly different from the one proposed by Zal & Gascoigne (54), although the actual numerical difference will be negligible for *l* _*A*_ << 1 and *d*_*D*_ << 1, which is generally the case.

### A.5 Advanced mixture case: *D, A* and {*D A*_*i*_ }_*i*=1…*n*_ mixture

In the more general case where a number *n* of distinct FRET configurations {*D A*_*i*_}_*i*=1…*n*_ of the donor and acceptor molecules can be observed (a similar derivation can be performed assuming a continuous probability distribution function rather than a finite number of species), with FRET efficiencies {*E*_*i*_} and fractions { *f*_*i*_}, the total number *N*_*D*_ of donor molecules and *N*_*A*_ of acceptor molecules can be decomposed into:

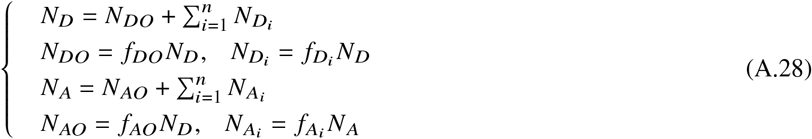

where 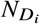 (resp.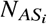) is the number of donor (resp. acceptor) molecules involved in a DA pair characterized by FRET efficiency *E*_*i*_, and *N*_*DO*_ (resp. *N*_*AO*_) is the number of remaining donor (resp. acceptor) molecules. Noting 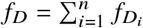 and 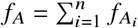, if follows from Eq. A.28 that *f*_*DO*_ = 1 − *f*_*D*_ and *f*_*AO*_ = 1 − *f*_*A*_.

By definition, the number of FRET pairs with FRET efficiency *E*_*i*_ is 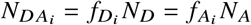. For each FRET pair, Eqs. A.9-A.11 apply and can be written (using the notations of Eq. A.14 modified to use *E*_*i*_ instead of *E* and *F*_*i*_ instead of *F*) as:

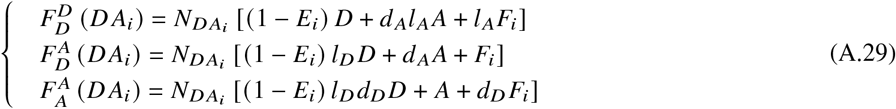

Eqs. A.23 for the mixture are now replaced by modified equations involving the multiple FRET species. We will only walk through the first equation, the last two following accordingly:

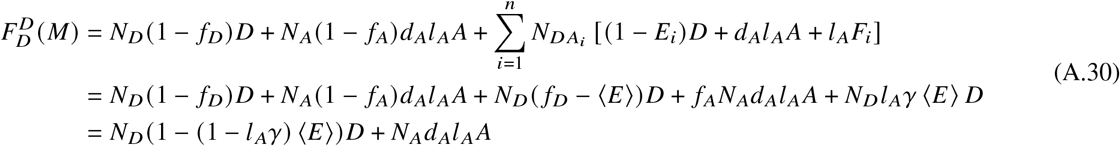

which is identical to the first equation in A.25 with the replacement of *f*_*D*_ *E* by ⟨*E*⟩ defined by:

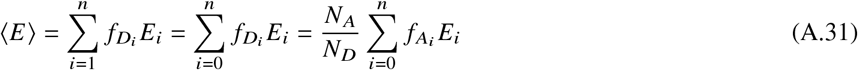

Following the same procedure for the remaining two equations for 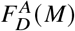 and 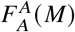 leads to the same equations as in A.25, with the replacement of *f*_*D*_ *E* by ⟨*E*⟩, Consequently, the result of Eq. A.27 applies, with the same replacement:

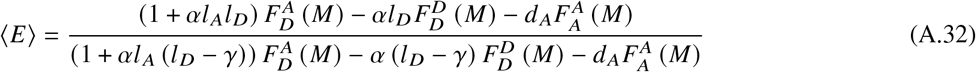

This equation allows comparing the results of intensity FRET measurements to those obtained by fluorescence lifetime in the case of multiple FRET efficiencies involving the same donor-acceptor pair (Eq. 17), with the identification 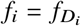.

