## Supplemental Note, Tables & Figures for "*in vivo* quantitative FRET small animal imaging: intensity versus lifetime-based FRET"

### SUPPLEMENTAL MATERIAL FOR THE ARTICLE "*IN VIVO* QUANTITATIVE FRET SMALL ANIMAL IMAGING: INTENSITY VERSUS LIFETIME-BASED FRET" BY SMITH *ET AL.*

#### SUPPLEMENTAL TABLES

Table S1: MFLI acquisition settings. For the *in vivo* MFLI measurements, illumination was adjusted [0 - 100%] over each organ (urinary bladder and liver) to prevent saturation. These changes are indicated for each organ as *frame:percentage*, where *frame* indicates the dataset number in the series, and *percentage* is the DMD reflection level (default: 100%).

| Experiment | MCP voltage | integration per gate image | Number of Gates | DMD Adjustment |
|---|---|---|---|---|
| *in vitro* IRF | 450 V | 100 ms | 176 | - |
| *in vitro* Fluorescence | 550 V | 150 ms | 176 | - |
| *in vivo* IRF | - | - | 151 | - |
| *in vivo* Fluorescence | 550 V | 150 ms | 161 | - |
| Urinary Bladder | - | - | - | 014:60% |
| Liver | - | - | - | - |

Table S2: IVIS exposure times

| Experiment | $F_{D_{ex}}^{D_{em}}$ | $F_{D_{ex}}^{A_{em}}$ | $F_{A_{ex}}^{A_{em}}$ |
|---|---|---|---|
| dsDNA | 1 s | 3 s | 2 s |
| Tf-TfR | 2 s | 2 s | 2 s |

Table S3: Intensity FRET correction parameters obtained using the dsDNA standards (Fig. 1).

| $l_D$ | $d_D$ | $l_A$ | $d_A$ |
|---|---|---|---|
| 0.28 ± 0.06 | 0.39 ± 0.03 | $1.5\times10^{-2} \pm 0.4\times10^{-2}$ | 0.43 ± 0.05 |

Table S4: $f_D E$ quantification obtained using both intensity and FLI-FRET for the dsDNA standards (Fig. 1).

| Technique | $DO$ | $DA_{27}$ | $DA_{22}$ | $DA_{17}$ |
|---|---|---|---|---|
| Intensity FRET | 3.0 ± 9.1 % | 18.0 ± 9.2 % | 22.8 ± 9.2 % | 50.4 ± 6.8 % |
| FLI-FRET | 2.1 ± 1.6 % | 24.7 ± 2.3 % | 33.3 ± 2.3 % | 52.5 ± 2.0 % |

Table S5: Measurements required for *in vivo* FRET analysis. Intensity FRET quantification required multiple animals, as well as multiple acquisition channels. MFLI-FRET only requires a single animal sample and single acquisition channel. (*) Indicates measurements that need to be repeated assuming equivalent experimental conditions. (**) When an internal negative FRET control is not available a donor-only labeled animal may be needed for *in vivo* lifetime FRET analysis.

| Intensity FRET | | | Lifetime FRET | | |
|---|---|---|---|---|---|
| No. | Sample | Channel | No. | Sample | Channel |
| 1 | Donor Injection | Donor | 1 | FRET injection* | Donor |
| 2 | Donor Injection | Acceptor | 2 | Donor injection** | Donor |
| 3 | Donor Injection | FRET | | | |
| 4 | Acceptor Injection | Donor | | | |
| 5 | Acceptor Injection | Acceptor | | | |
| 6 | Acceptor Injection | FRET | | | |
| 7 | FRET Injection* | Donor | | | |
| 8 | FRET Injection* | Acceptor | | | |
| 9 | FRET Injection* | FRET | | | |

#### SUPPLEMENTAL FIGURES

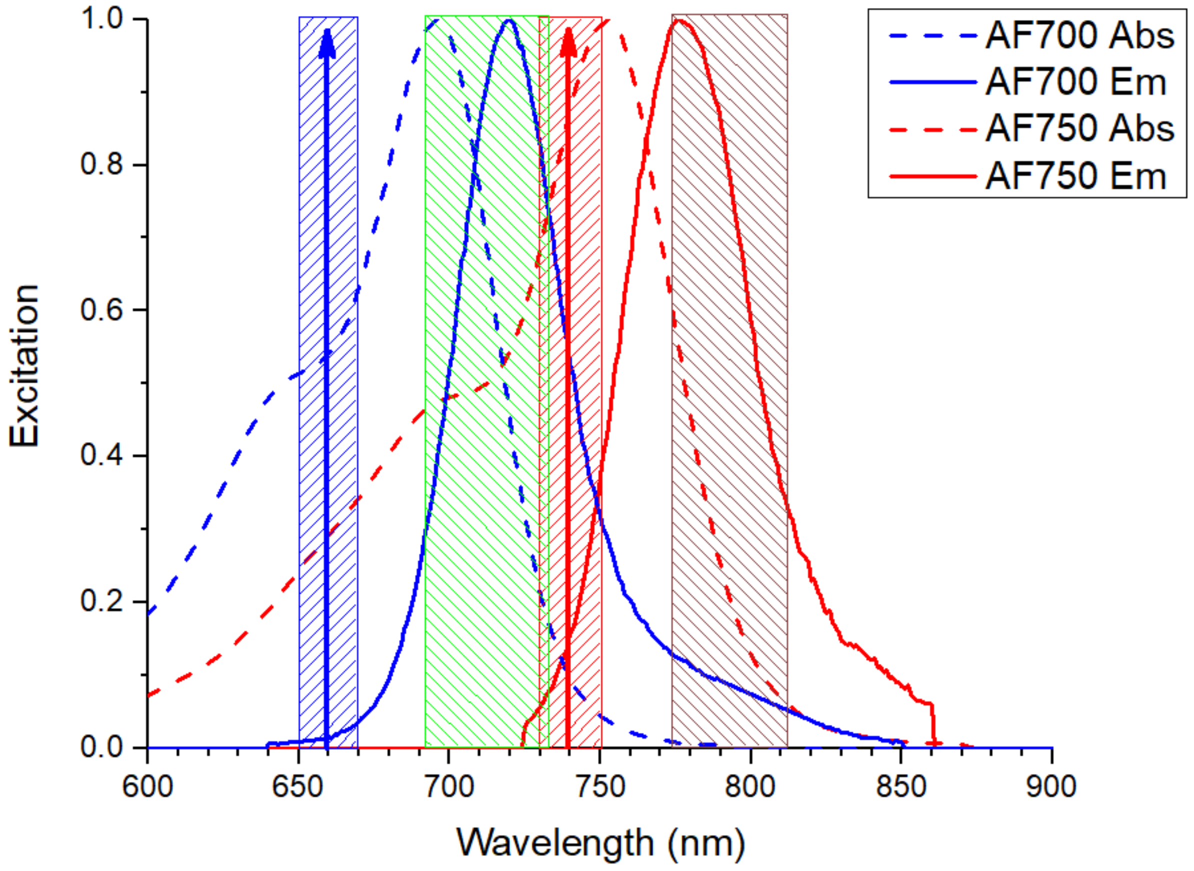


Figure S1: Excitation (dashed curves) and emission spectra (plain curves) of AF700 and AF750, as well as the excitation and emission filter characteristics used in the intensity-based measurements (hashed stripes, the vertical arrows indicating the center of the excitation filters).

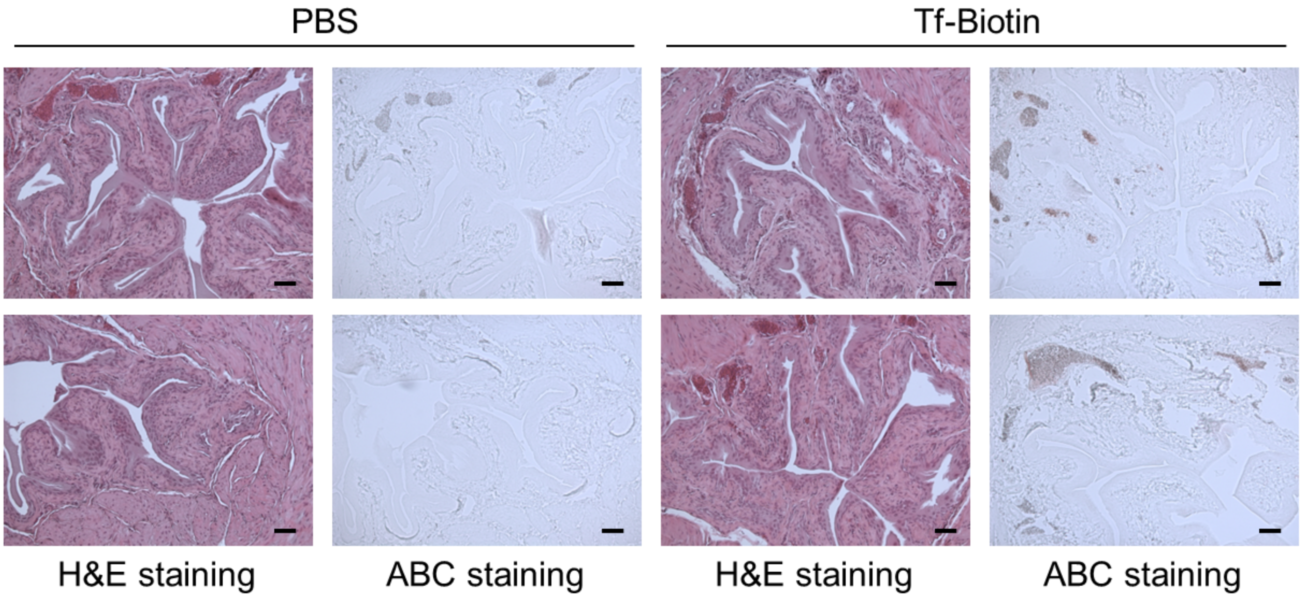


Figure S2: Two mice were injected with biotin-Tf or PBS and sacrificed 6 hr post-injection. Bladders were collected and processed for immunohistochemistry using ABC Elite and NovaRed kit (ABC staining) to visualize Tf-biotin. Parallel bladder sections were stained with Hematoxylin and Eosin (H&E staining) and imaged using a 10x microscope for tissue morphology visualization. H&E staining shows urinary cavity surrounded by papillary transitional, epithelium and thick wall of smooth muscle and fat cells. ABC staining shows weak positive staining in fat cells, consistent with reduced FRET signal in urinary bladder. Scale bar: 100 µm.

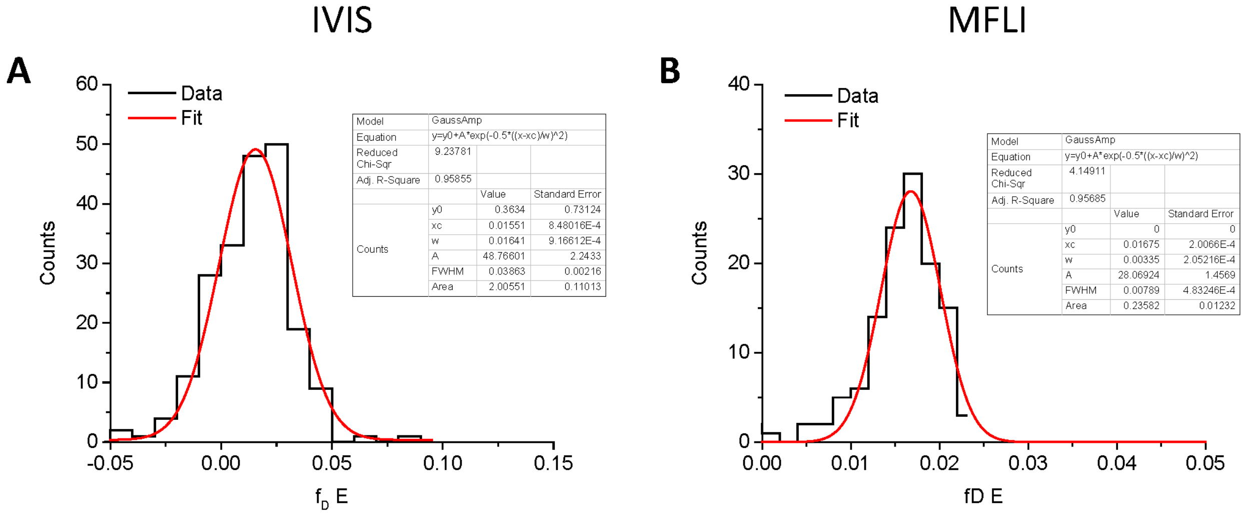


Figure S3: Histograms and normal fits of $f_D E$ values measured in the urinary bladders of both mice over the duration of the measurement (see Fig. 5B & D). A: IVIS measurements ($f_D E$ = 1.6 ± 1.6%). B: MFLI measurements ($f_D E$ = 1.7 ± 0.3%).

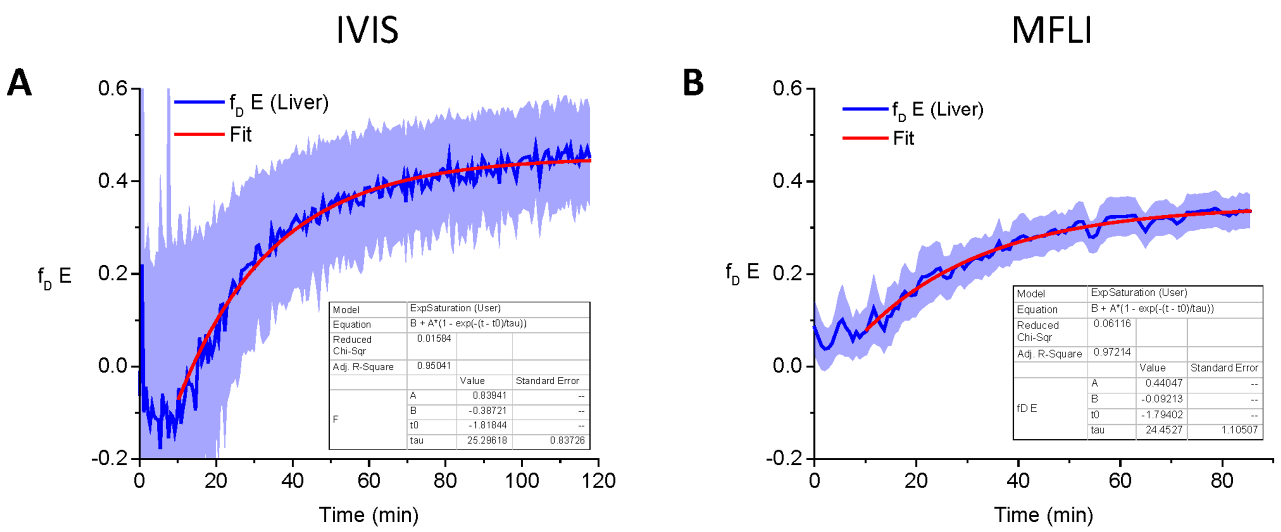


Figure S4: Pharmaco-kinetic time constants of Tf-TfR engagement in the mouse liver measured using intensity-FRET or lifetime-FRET methods. The $f_D E$ curves shown in Fig. 5B & D were fitted with a single-exponential model: $f_D E(t) = A\,(1 - \exp\left(-(t - t_0)/\tau_{kin}\right)) + B$ for t > 10 min (red curves), resulting in similar kinetic time constants: $\tau_{kin,IVIS}$ = 25.3 ± 0.8 min and $\tau_{kin,MFLI}$ = 24.5 ± 1.1 min (these are two different mice). Note that the $\tau_{kin,MFLI}$ value obtained here using single-pixel analysis is similar to that obtained previously at the whole organ level in ref. (46), Fig. S6 (mouse 2:1), where $\tau_{kin,MFLI}$ = 29.1 ± 1.1 min was computed. The asymptotic mean FRET efficiencies ($f_D E(t \to \infty) = A + B$) are 0.45 (IVIS) and 0.55 (MFLI).

#### S1 SUPPLEMENTAL NOTE: INTRINSIC FLUORESCENCE BACKGROUND AND DONOR-ONLY LIFETIME CHOICE IN MFLI

##### S1.1 Intrinsic fluorescence background

To study the influence of background autofluorescence on data analysis in the MFLI experiment discussed in this article, two mice injected only with Af700-Tf (donor) were studied with the same approach as described in the main text.

Importantly, the two mice were recorded for ≈10 min prior to probe injection, which allowed analyzing the intensity and temporal profile of the autofluorescence signal excited by the 695 nm, 100 fs pulsed laser excitation in nude mice and compare it to the signal observed after injection. The effect of this background fluorescence signal on lifetime analysis is discussed in section S1.2.

Fig. S5A-C shows snapshots of one of the mice observed for approximately 2 hours with the MFLI system, starting with a window of 12 time points (000-011) during which no fluorophore was present in the mouse (a test tube with AF700, visible as a bright spot on the left of panels A-C, was present next to the mouse as a control). Injection of AF700-Tf only resulted in the appearance of signal throughout the mouse (starting at time point 012, shown in panel B), with increasing intensity observed in the urinary bladder (UB) and liver as time went on. The "background" signal originally observed in the UB (black TPSF in panel D) and in the liver (black TPSF in panel E) is very different in shape and amplitude from the fluorescence signal recorded in the UB and the liver immediately after injection (time point 012, blue curves in panels D & E) or at the end of the recording (time point 164, red curves in panels D & E).

The source of this background signal is unclear and we therefore use the generic term of "tissue autofluorescence". It is unlikely to be due to two-photon excitation of NADH or FAD, whose emission is further in the blue-to-red region of the spectrum (74). Its lifetime is of the order of 0.5-0.6 ns, which is comparable to the lifetimes of interest for our MFLI-FRET studies, but in the conditions of our experiments, remains below 12% of the total signal (and for most of the time, in the low single digit percent of the total signal).

Background subtraction was not possible in the experimental case discussed in the main text, because the MFLI experiment started only after injection of donor- and acceptor-labeled Tf. Based on the repeated negative result illustrated in Figs. S5 & S6, obtained in similar conditions, we believe that that background does not affect our results, although it could potentially do so in the case of signals with lower intensities. It is therefore important to characterize it beforehand, or include a few frames prior to injection to use as reference and subtract from the remainder of the time series.

##### S1.2 Organ- and time-dependence of the donor-only lifetime

The donor-only lifetime $\tau_{DO}$ is critical to compute the mean FRET efficiency using Eq. 13. $\tau_{DO}$ is the donor-only lifetime of the isolate probe in a similar biochemical environment as that of other donor undergoing FRET with acceptor molecules.

To understand the dependence of the donor lifetime on the organ in which it accumulates, as well as its potential temporal variations, experiments with mice injected only with donor-labeled Tf molecules (AF700-Tf) were performed and analyzed using the same approach described in Material and Methods, but at the whole ROI level (instead of the single-pixel level used in the remainder of this study) to minimize the effect of low SNR. Fig. S6 shows that the donor lifetime is to some extent dependent on the individual mouse under study (labeled M0 and M5), although the two mice studied herein exhibit similar trends for the AF700-Tf (donor) lifetime in both the urinary bladder (UB, Fig. S6B & E) and the liver (Fig. S6C & F). The donor lifetime appears fairly constant in the UB after a brief initial decrease (M0 range: 1.06-1.18 ns, M5 range: 0.94-1.04 ns), while it exhibits a clear steady, if minimal, increase in the liver (M0 range: 0.95-1.09 ns, M5 range: 0.88-1.04 ns). In both cases, it is also noteworthy that these lifetimes are in general larger than that measured in a nearby test tube containing a concentrated solution of AF700 (M0: 1.0 ± 0.01 ns, M5: 0.91 ± 0.02 ns), possibly due to pH or concentration effects. It appears reasonable in these experiments to use the donor lifetime measured in the urinary bladder as that of the donor-only lifetime, as it is no different from that observed in mice injected with donor-only.

Finally, it is important to note that the background autofluorescence characterized in section S1.1 has little effect on the lifetime fitting analysis, as demonstrated in Fig. S6, which shows the fitted lifetimes in the two organs (UB & liver) with (+, red curves) or without (-, black curves) subtraction of the first time point (dataset 000).

##### S1.3 Choice of donor-only lifetime in *in vivo* MFLI-FRET analysis

The data discussed in the previous section shows that, to a good approximation, the donor-only (AF700-Tf) lifetimes in the UB and liver are similar. It is moreover noteworthy that, as shown in these measurements, the donor lifetime may depend not only on the location (e.g. urinary bladder or liver), but also on the time point of observation (i.e. shortly after injection or later on). These changes are however fairly minimal and the resulting uncertainty on the mean FRET efficiency (coming from the factor

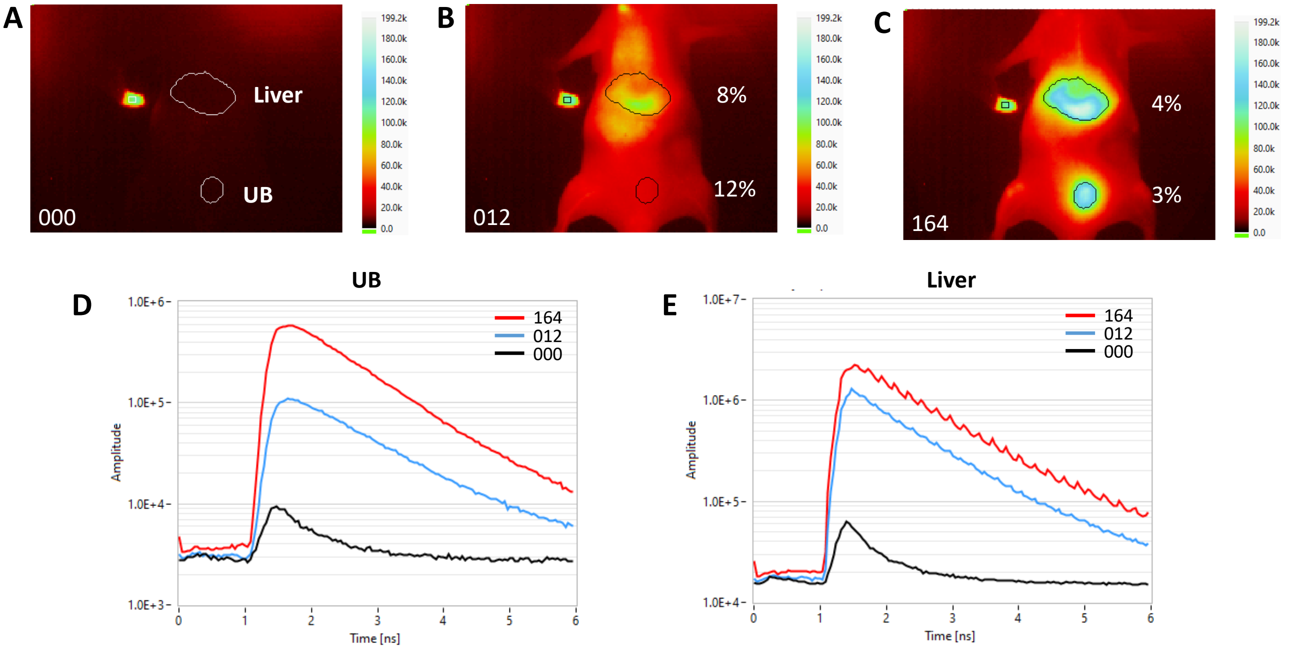


Figure S5: The background signal (taken as the signal observed at the first time point shown in A) represents only a fraction of the observed signal (indicated in white characters next to each organ in B & C), even at the first time point where fluorescence is observed (B), or at the last time point (where the signal is maximal, C). Comparison of background TPSFs (in black in D & E) for the UB and liver ROIs, to the signal observed at different time points in these same organs (blue: first time point with fluorescence, corresponding to B, red: last time point, corresponding to C).

$1/\tau_{DO}$ in Eq. 13) would be of the same order of magnitude.

From an experimental point of view, because these donor-only lifetime measurements are not possible during a FRET experiment (the best that could be achieved would be to first inject the donor and characterize its lifetime, and only then inject the acceptor, as performed in ref. (46)), the only practical solution is to use either:

- the largest lifetime component in the fit as that of the donor,
- a valid internal reference measured in a different region of the mouse (*e.g.* the urinary bladder).

The first option is illustrated in Fig. S7 for the mouse dataset discussed in Fig. 5C-D of the main text. Analysis was done at the ROI level to minimize the effect of low SNR and speed up analysis (*i.e.* the TPSFs of all pixels in the ROI were summed and analyzed as a single TPSF).

Panel A shows the evolution of the 4 parameters ($A_0$, $\tau_0$, $A_1$ and$\tau_1$) of a 2-Exp tail fit of the liver TPSF, when not constraining any of the parameters. The amplitude-averaged lifetime ($\langle\tau\rangle_a$, blue curve) is a derived parameter computed according to Eq. 10 of the main text. All parameters exhibit large fluctuations, in particular during the first half of the time series (notice a number of $\tau_1 = 0$ values and corresponding small $\langle\tau\rangle_a$ values), indicating that some of the fit parameters may need to be constrained (no such fluctuations are expected biophysically).

The distribution of each lifetime parameter is approximately normal with $\tau_0 = 1.10 \pm 0.08$ ns and $\tau_1 = 0.27 \pm 0.06$ ns (Fig. S8A), but can be modelled by a quasi-exponential decay of parameter $\tau_0$ to an asymptotic value $\tau_0 \approx 1.03$ ns and $\tau_1 \approx 0.21$ ns (Fig. S8B). The other parameters undergo similar steady changes, occasionally interrupted by non-physical jumps. To avoid these fluctuations due to fit instabilities, it is possible to progressively constrain the two lifetimes, interpreted as that of the donor-only ($\tau_0$) and that of the FRET-undergoing donor ($\tau_1$) around their mean values. This results in the curves shown in Panel B of Fig. S7 (both lifetimes constrained to stay within $3\sigma$ of their mean values), Panel C (both lifetimes constrained to stay within $2\sigma$ of their mean values), Panel D (both lifetimes constrained to stay within $1\sigma$ of their mean values) and finally Panel E (both lifetimes constrained to be equal to their mean values).

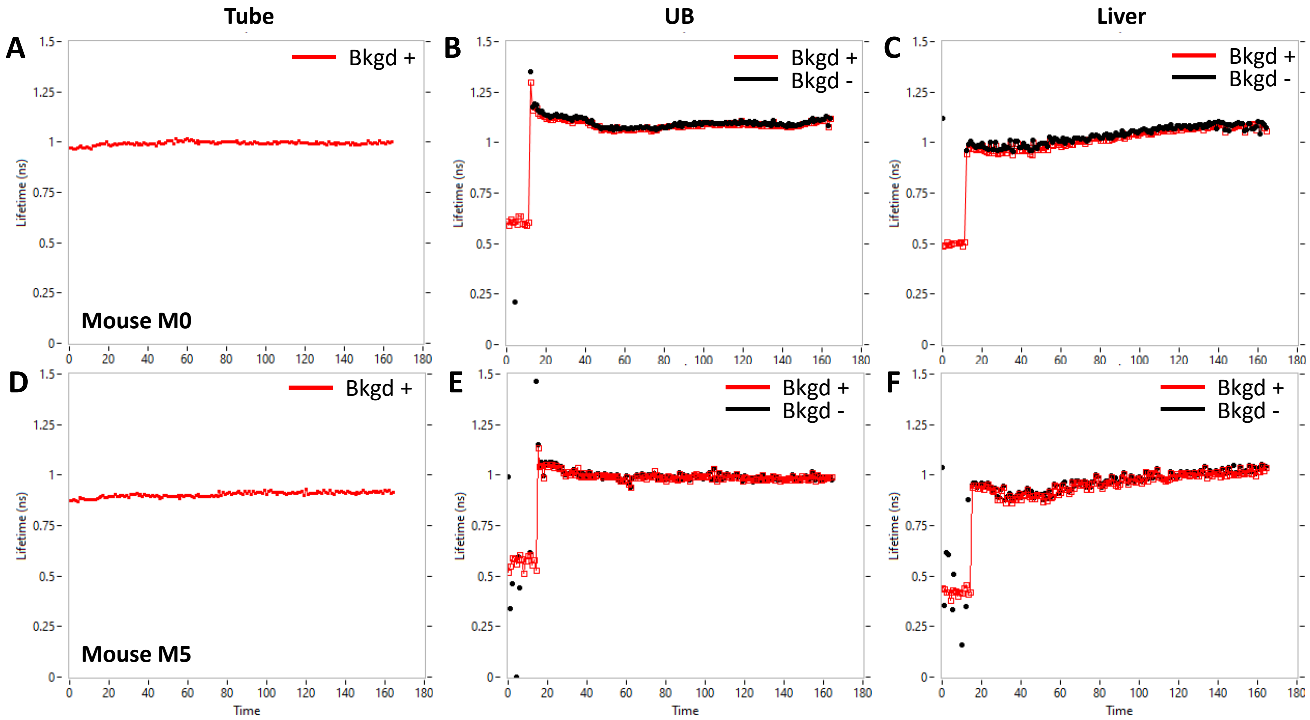


Figure S6: Fitted ROI lifetimes with (-, red) or w/o (+, black) background subtraction, showing that there is no significant effect of background on the fitted lifetimes. (A-C): Mouse M0, (D-F): Mouse M5. There are however noticeable changes of the measured lifetime over time in both organs in the two mice. The *Time* axis indicate the frame number (0 to 164), each frame being separated from the next by ≈ 43 s.

As shown in Panel F, which reproduces the different derived parameters $\langle\tau\rangle_a$ obtained from these different analyses, they all return the same result within a few percent. In other words, despite the constraints imposed on the two lifetimes, the general conclusion gleaned from a qualitative look at Panel A is not modified: there is progressive decrease of the mean lifetime. Constraining the component naturally interpreted as the donor-only lifetime does hide the slight decrease of its value observed in the absence of constraint (Fig. S7A & S8B), but this decrease is relatively small ($\Delta\tau_0 \approx 0.22$ ns, of the order of the range of the donor lifetime fluctuations observed in Fig. S6 for the donor-only injected mice). This variation could be due to a variety of reasons, including changes in the electronic environment of the donor-labeled probes, or the appearance of a weak FRET interaction between donor and acceptor brought in proximity by some unspecified mechanism.

In the absence of additional information to elucidate the origin of this variation, and because it does not affect the overall conclusion of the study (Fig. S7F), this analysis indicates that the approach consisting in using the longest fitted lifetime for the donor only is justified. However, because of the lower signal available when performed at the single-pixel level, this approach will result in a lot more noise, which is why constraints of the type used in Panel B-E are generally used to avoid excessive (and not biophysical) fluctuations.

The second option of using an independent (and internal) reference for the donor lifetime is illustrated in Fig. S8. As indicated by the results of Fig. S6 (donor-only injected mice), the liver donor lifetime should be close to that observed in the urinary bladder, and fairly (but not exactly) constant. As shown in Fig. S8D, the mouse studied here is characterized by a very stable urinary bladder donor lifetime (red curve, $\tau_0 = 1.01 \pm 0.01$ ns), which we can use as a constraint for the longer lifetime of the 2-Exp fit. The result is illustrated in Fig. S8B, which still shows large fluctuations of the second lifetime component (black dots) and its amplitude (open black squares). However, it is clear that these fluctuations stabilize around time point 40 to a value $\tau_1 = 0.22 \pm 0.02$ ns. This lifetime is again naturally interpreted as the quenched donor-lifetime (or possibly the average of many distinct quenched donor lifetimes), which there is no reason to suspect to be any different at the end of the observation from what it would be at the beginning (FRET occurring between an AF700-Tf bound to a dimeric TfR receptor and an AF750-Tf bound to the same receptor). However, it is expected that its fraction (or equivalently, its amplitude) might increase over time, as there is indication of happening in Fig. S8A past the initial fluctuations (open black squares). As argued above when discussing the use of the longest lifetime component for the donor lifetime, it is natural to attempt to avoid these non-physical fluctuations by imposing constraints on the second lifetime parameter. For instance, by setting $\tau_1$ = 0.22 ns and only fitting the unknown amplitudes of each components. The result is shown in Fig. S8C, which demonstrates a steady increase of the quenched-donor amplitude (open black squares) with a correlated decrease of the free donor amplitude (open red squares), resulting in a monotonous decrease of the average lifetime $\langle \tau \rangle_a$ (small blue squares). The resulting $\langle \tau \rangle_a$ curve is reproduced in Fig. S8D next to the UB lifetime for comparison.

Comparison of the inferred average FRET efficiency $\langle E \rangle$ (Eq. 11, open red circles) obtained by this approach and the corresponding $\langle E \rangle$ obtained with the longest lifetime reference approach (open blue squares) is shown in Fig. S8E. The single-pixel analysis presented in Fig. 5D of the main text is also reproduced as a black curve. While there are some small quantitative differences between the different approaches, the same general conclusion, *i.e.* that an increasing amount of FRET is observed in the liver, is obtained in all cases.

In summary, while it is obvious that obtaining a precise value of the pure donor (AF700-Tf) lifetime in the liver is challenging in the presence of FRET, the prior knowledge of its minimal variation over the course of such an experiment, as well as its similarity to that observed in the urinary bladder, allows to confidently use either approach to interpret the data. While we cannot exclude longer distance donor-acceptor interactions resulting in an observed decay component slightly shorter than the pure donor lifetime (potentially the reason for the apparent decrease of the long lifetime component observed in Fig. S8B), this variation is small enough to not affect the general conclusion of these experiments, namely the progressive increase of the average FRET efficiency $\langle E \rangle$ in the liver over time. This approximation may not be applicable in other applications.

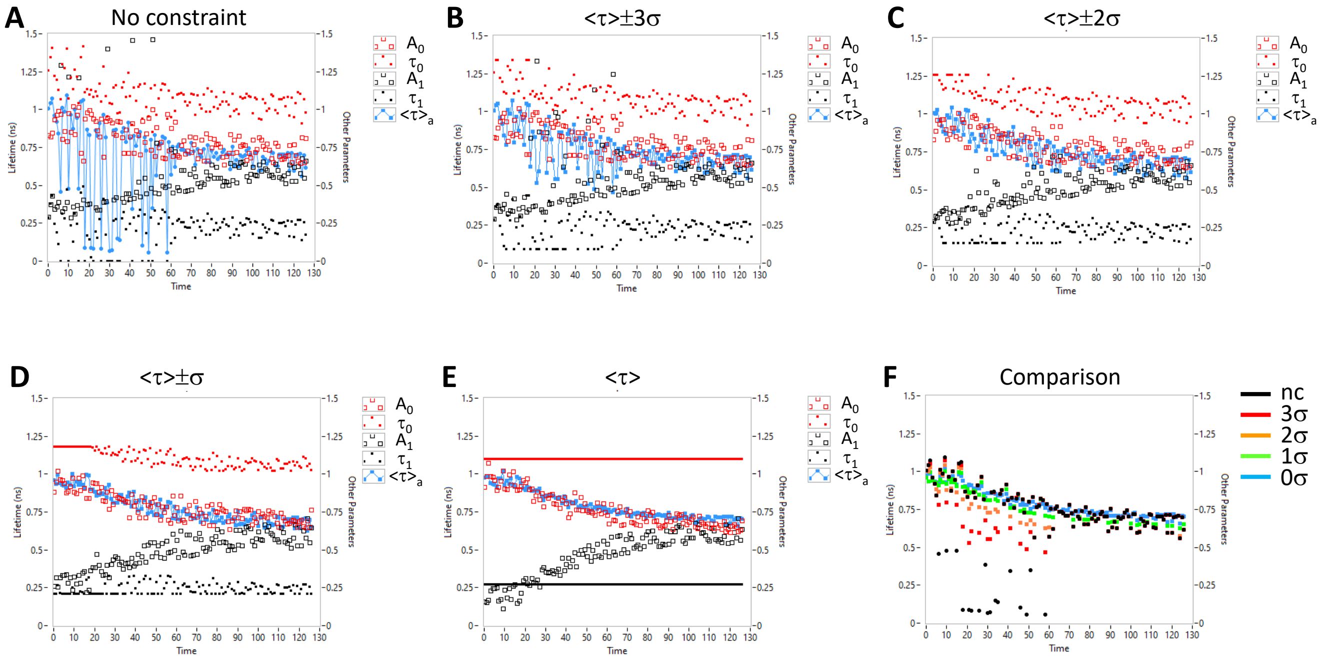


Figure S7: Whole liver ROI FLI analysis with different lifetime parameter constraints. A: $\tau_i \in [0, 5]$, B: $\tau_i \in [< \tau_i > -3\sigma, < \tau_i > +3\sigma]$, C: $\tau_i \in [< \tau_i > -2\sigma, < \tau_i > +2\sigma]$, D: $\tau_i \in [< \tau_i > -\sigma, < \tau_i > +\sigma]$, E: $\tau_i =< \tau_i >$. F: Comparison of the computed $< \tau >_a$ curves for the different conditions. The *Time* axis indicate the frame number (1 to 127) minus 1, each frame being separated from the next by 41 s.

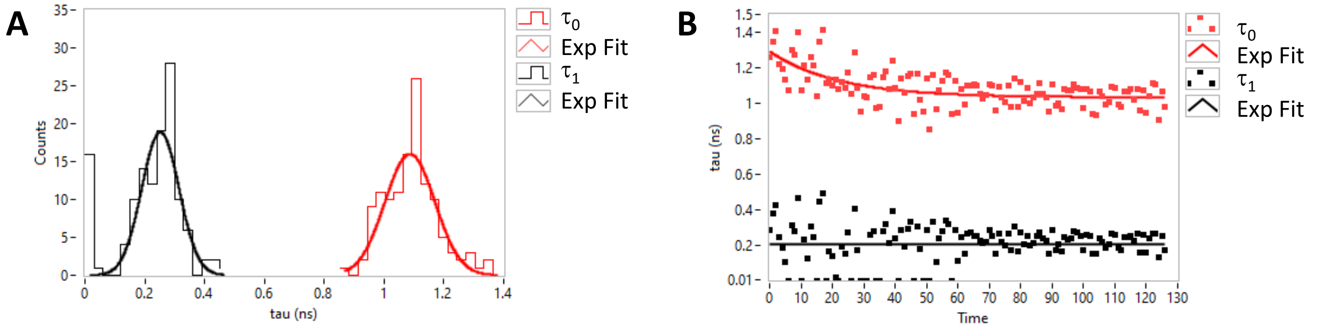


Figure S8: A: Histograms of fitted $\tau_0$ and $\tau_1$ (unconstrained parameters) and Gaussian fits. B: Same parameters represented as a function of time frame. Single-exponential fits of these distribution are shown as plain curves. The *Time* axis indicate the frame number (1 to 127) minus 1, each frame being separated from the next by 41 s.

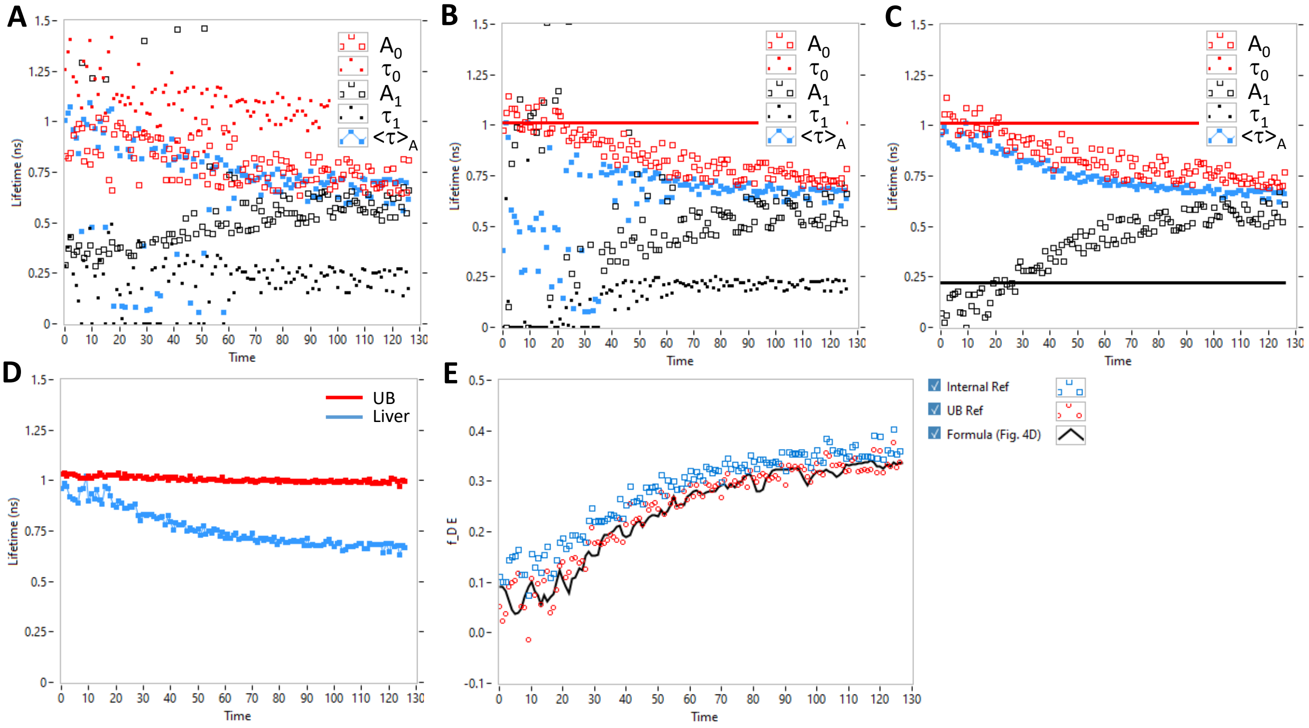


Figure S9: ROI-level FRET analysis. A-C: Liver ROI NLSF analysis with a 2-Exp model limited to 2-99% of the tail. A: with no constraints on $\tau_0$ & $\tau_1$ (identical to Panel A of Fig. S7), B: with fixed $\tau_0$ = 1.01 ns and C: with $\tau_0$ = 1.01 ns & $\tau_1$ = 0.22 ns. D: Comparison of the average donor lifetime in the urinary bladder (UB, red) and in the liver (blue, result from B). E: Comparison of this ROI-level analysis (blue) to the single-pixel analysis described in the main text (black, from Fig. 5D). The *Time* axis indicate the frame number (1 to 127) minus 1, each frame being separated from the next by 41 s.
